# Musical expertise shapes functional and structural brain networks independent of absolute pitch ability

**DOI:** 10.1101/2020.07.24.216986

**Authors:** Simon Leipold, Carina Klein, Lutz Jäncke

## Abstract

Professional musicians are a popular model for investigating experience-dependent plasticity in human large-scale brain networks. A minority of musicians possess absolute pitch, the ability to name a tone without reference. The study of absolute pitch musicians provides insights into how a very specific talent is reflected in brain networks. Previous studies of the effects of musicianship and absolute pitch on large-scale brain networks have yielded highly heterogeneous findings regarding the localization and direction of the effects. This heterogeneity was likely influenced by small samples and vastly different methodological approaches. Here, we conducted a comprehensive multimodal assessment of effects of musicianship and absolute pitch on intrinsic functional and structural connectivity using a variety of commonly employed and state-of-the-art multivariate methods in the largest sample to date (n = 153 female and male human participants; 52 absolute pitch musicians, 51 non-absolute pitch musicians, and 50 non-musicians). Our results show robust effects of musicianship in inter- and intrahemispheric connectivity in both structural and functional networks. Crucially, most of the effects were replicable in both musicians with and without absolute pitch when compared to non-musicians. However, we did not find evidence for an effect of absolute pitch on intrinsic functional or structural connectivity in our data: The two musician groups showed strikingly similar networks across all analyses. Our results suggest that long-term musical training is associated with robust changes in large-scale brain networks. The effects of absolute pitch on neural networks might be subtle, requiring very large samples or task-based experiments to be detected.

**Significance Statement:** A question that has fascinated neuroscientists, psychologists, and musicologists for a long time is how musicianship and absolute pitch, the rare talent to name a tone without reference, are reflected in large-scale networks of the human brain. Much is still unknown as previous studies have reported widely inconsistent results based on small samples. Here, we investigate the largest sample of musicians and non-musicians to date (n = 153) using a multitude of established and novel analysis methods. Results provide evidence for robust effects of musicianship on functional and structural networks that were replicable in two separate groups of musicians and independent of absolute pitch ability.

## Introduction

Professional musicians are a commonly studied model for experience-dependent brain plasticity (Münte et al., 2002; Jäncke, 2009; Schlaug, 2015). Intense musical training starting early in life is thought to cause neuroplastic adaptations that are paralleled by improvements in audition, sensory-motor skills, and possibly higher-order cognitive functions (Fujioka et al., 2006; Hyde et al., 2009; Seither-Preisler et al., 2014; Habibi et al., 2018). In recent years, a major focus within the neuroscience of music has been on training-related plasticity in large-scale brain networks, which underlie most human sensory, motor, and cognitive functions (Bressler and Menon, 2010).

Previous research provides evidence that musicianship is associated with differences in both the intrinsic functional and structural networks of the human brain. However, an examination of these studies reveals inconsistencies in findings regarding the location of the effects in the brain and also the direction of these effects. For example, while most of the studies report hyperconnectivity in musicians compared to nonmusicians (Fauvel et al., 2014; Klein et al., 2016), others have found hypoconnectivity (Imfeld et al., 2009), or both (Schmithorst and Wilke, 2002; Bengtsson et al., 2005). These studies suggest that in musicians, connectivity between brain regions is altered across the entire brain including sensory, motor, multisensory, and cognitive regions of the cortex (Klein et al., 2016; Palomar-García et al., 2017), subcortex (Luo et al., 2012), and the cerebellum (Abdul-Kareem et al., 2011).

The diversity of these findings could be influenced by small sample sizes and inconsistent methodology. In studies examining intrinsic functional connectivity, the number of participants in the musician groups ranged from 10 (Zamorano et al., 2017) to 25 (Luo et al., 2014), and in studies examining structural connectivity, from only five (Schmithorst and Wilke, 2002) to 36 (Steele et al., 2013). Studies with small samples lack the statistical power to detect small effects, and findings from small-scale studies have a higher probability of returning false positives (Button et al., 2013). With regard to methodology, many previous studies took a region of interest (ROI)-based approach. To our knowledge, only two functional connectivity studies exist using a data-driven, connectomic whole-brain approach (Luo et al., 2014; Klein et al., 2016). Studies on structural networks in musicians have exclusively used an ROI-based approach by focusing on separate white-matter tracts or brain regions. No previous structural connectivity study comparing musicians and non-musicians has employed a whole-brain connectomic approach.

Apart from general effects of musicianship, some studies have focused on a special talent present among musicians: absolute pitch (AP), the rare ability to name a tone without reference (Deutsch, 2013). Only a few studies examined intrinsic functional networks in AP versus non-AP musicians. Again, the findings of these studies show little consistency, suggesting an effect of AP on functional connectivity of sensory, parietal, and frontal cortex (Elmer et al., 2015; Kim and Knösche, 2017; Brauchli et al., 2019). The applied methodology differed widely between studies (cf. Jäncke et al., 2012; Loui et al., 2012; Wenhart et al., 2019). An effect of AP on structural connectivity has been reported in the vicinity of associative auditory areas (Loui et al., 2011; Dohn et al., 2015; Kim and Knösche, 2016; Burkhard et al., 2020). None of the previous studies investigating AP and structural connectivity employed a whole-brain connectomic approach. Importantly, all of these results have yet to be replicated in an independent sample.

Taken together, findings from previous studies are highly inconsistent, possibly due to small samples and methodological differences. In this study, we aimed to identify robust effects of musicianship and AP on functional and structural connectivity using a multitude of previously employed and novel methods on a large multimodal dataset (n = 153), consisting of 52 AP musicians, 51 non-AP musicians, and 50 nonmusicians. We employed ROI-based and whole-brain approaches, and a multivariate approach based on machine learning algorithms. Crucially, we determined if effects of musicianship were replicable in both musician groups, irrespective of their AP ability.

## Materials and methods

### Participants

We analyzed resting-state functional magnetic resonance imaging (rsfMRI) and diffusion-weighted imaging (DWI) data of 153 female and male human participants. A portion of the rsfMRI data (Brauchli et al., 2019) and the DWI data (Burkhard et al., 2020) was previously analyzed using a different methodology. The participants consisted of three groups: AP musicians (n = 52), non-AP musicians (n = 51), and nonmusicians (n = 50). The groups were comparable regarding sex, handedness, age, rsfMRI movement, and DWI movement (see Table 1). Participants of the musician groups were either professional musicians, music students, or highly trained amateurs. Assignment to the musician groups (AP or non-AP) was based on self-report and confirmed by a tone-naming test (Oechslin et al., 2010a, 2010b). During the test, participants had to name 108 pure tones presented in a pseudorandomized order. Octave errors were disregarded in the calculation of the tone-naming score. In the rare case that a potential participant had indicated to be an AP musician in the initial online application form but then performed around chance level (8.3%) in the tone-naming test, we did not invite this individual to undergo the imaging experiments in the laboratory. In contrast, we did invite individuals who had indicated to be non-AP musicians and then showed a high level of proficiency in tone-naming that was above chance level (and reiterated in the laboratory that they do not possess AP); we did not regroup these participants as AP musicians (Leipold et al., 2019a). Non-musicians had not received formal musical training in the five years prior to the study.

**Table 1.** Participant characteristics.

|  | **AP musicians** | **Non-AP musicians** | **Non-musicians** |
| --- | --- | --- | --- |
| Number of participants | 52 | 51 | 50 |
| Sex (female / male) | 24 / 28 | 24 / 27 | 24 / 26 |
| Handedness (right / left / both) | 45 / 4 / 3 | 46 / 4 / 1 | 44 / 6 / 0 |
| Age | 26.37 ± 4.98 years | 25.29 ± 4.42 years | 25.86 ± 5.52 years |
| rsfMRI movement <sup>+</sup> | 8.90 ± 16.31 scans | 5.61 ± 11.77 scans | 5.26 ± 15.43 scans |
| DWI movement <sup>§</sup> | 0.47 ± 0.11 | 0.48 ± 0.11 | 0.44 ± 0.12 |
| Musical aptitude (AMMA) – total | 66.04 ± 6.18 | 63.45 ± 6.96 | 52.80 ± 9.22 |
| Musical aptitude (AMMA) – tonal | 32.33 ± 3.67 | 30.55 ± 4.23 | 25.34 ± 5.02 |
| Musical aptitude (AMMA) – rhythm | 33.71 ± 2.78 | 32.90 ± 3.03 | 27.46 ± 4.58 |
| Tone-naming score | 76.41 ± 19.96 % | 23.66 ± 19.16 % | 8.41 ± 3.52 % |
| Age of onset of musical training | 6.06 ± 2.40 | 6.53 ± 2.39 | – |
| Years of musical training | 20.31 ± 5.26 years | 18.76 ± 5.01 | – |
| Cumulative musical training | 16,347.68 ± 12,582.35 hours | 13,830.10 ± 9,985.04 hours | – |

Abbreviations: AMMA = Advanced Measures of Music Audiation; AP = absolute pitch; DWI = diffusion-weighted imaging; FD = framewise displacement; rsfMRI = resting-state functional magnetic resonance imaging.

Demographical (sex, handedness, age) and behavioral data (musical aptitude, musical experience, and tone-naming proficiency) were collected using LimeSurvey (https://www.limesurvey.org/). Self-reported handedness was confirmed using a German translation of the Annett questionnaire (Annett, 1970). Musical aptitude was assessed using the Advanced Measures of Music Audiation (AMMA) (Gordon, 1989). During the AMMA test, participants were presented with short pairs of piano sequences. The participants had to decide whether the sequences were equivalent or differed in tonality or rhythm. None of the participants reported any neurological, audiological, or severe psychiatric disorders, substance abuse, or other contraindications for MRI. All participants provided written informed consent and were paid for their participation or received course credit. The study was approved by the local ethics committee (https://kek.zh.ch/) and conducted according to the principles defined in the Declaration of Helsinki.

### Experimental design and statistical analysis

#### Statistical analysis of behavioral data

Participant characteristics were compared between the groups using one-way analyses of variance (ANOVAs) with a between-participant factor *group* or Welch’s *t*-tests where appropriate (significance level α = 0.05). The analyses were performed in R (version 3.6.0, RRID:SCR_001905). We used the R packages *ez* (version 4.4-0, https://CRAN.R-project.org/package=ez) for frequentist ANOVAs and *BayesFactor* (version 0.9.12-4.2, https://CRAN.R-project.org/package=BayesFactor) for Bayesian ANOVAs (Rouder et al., 2012) and Bayesian *t*-tests (Rouder et al., 2009). We used default priors as implemented in the *BayesFactor* package. Consequently, alongside *p* values, we report Bayes factors quantifying the evidence for the alternative relative to the null hypothesis (BF10) and vice versa (BF01). Bayes factors are interpreted as evidence for one hypothesis relative to the other hypothesis. A Bayes factor between 1 and 3 is considered as anecdotal evidence, between 3 and 10 as moderate evidence, between 10 and 30 as strong evidence, between 30 and 100 as very strong evidence, and larger than 100 as extreme evidence. Effect sizes of ANOVA-effects are given as generalized eta-squared (η^2^_G_) and effect sizes for *t*-tests are given as Cohen’s *d.*

#### MRI data acquisition

Magnetic resonance imaging (MRI) data were acquired using a Philips Ingenia 3.0T MRI system (Philips Medical Systems, Best, The Netherlands) equipped with a commercial 15-channel head coil. For each participant, we acquired whole-brain rsfMRI and DWI data, and a whole-brain anatomical T1-weighted image to facilitate the spatial normalization of the rsfMRI and DWI data. For the musician groups, we also collected fMRI data during a pitch-processing task, which is discussed in another publication (Leipold et al., 2019a). The whole scanning session lasted around 50 minutes.

#### rsfMRI data acquisition

For the acquisition of rsfMRI data, we used a T2*-weighted gradient echo (GRE) echo-planar imaging (EPI) sequence with the following parameters: repetition time (TR) = 2,300 ms, echo time (TE) = 30 ms, flip angle α = 78°, slice scan order = interleaved, number of axial slices = 40, slice thickness = 3 mm, field of view (FOV) = 220 x 220 x 143 mm^3^, acquisition voxel size = 3 x 3 x 3 mm^3^; reconstructed to a spatial resolution of 2.75 x 2.75 x 3.00 mm^3^ with a reconstruction matrix of 80 x 80, number of dummy scans = 5, total number of scans = 210, total scan duration = 8 min. Participants were instructed to relax and look at a fixation cross during the scanning.

#### DWI data acquisition

We acquired DWI data using a diffusion-weighted spin echo (SE) EPI sequence with the following parameters: TR = 10,022 ms, TE = 89 ms, acquisition and reconstructed voxel size = 2 x 2 x 2 mm^3^, reconstruction matrix = 112 x 112, flip angle α = 90°, FOV = 224 x 224 x 152 mm^3^, number of axial slices = 76, B = 1000 s/mm^2^, number of diffusion-weighted scans/directions = 64, number of non-diffusion weighted scans = 1, total scan duration = 14 min. Additionally, we acquired six non-diffusion weighted images (B = 0) in opposing phase-encoding directions (anterior-posterior, posterior-anterior), which were used during the preprocessing of the DWI data.

#### T1-weighted MRI data acquisition

The anatomical image was acquired using a T1-weighted GRE turbo field echo sequence with the following parameters: TR = 8.1 ms, TE = 3.7 ms, flip angle α = 8°, number of sagittal slices = 160, FOV = 240 x 240 x 160 mm^3^, acquisition voxel size = 1 x 1 x 1 mm^3^; reconstructed to a spatial resolution of 0.94 x 0.94 x 1.00 mm^3^ with a reconstruction matrix of 256 x 256, total scan duration = 6 min.

#### MRI data preprocessing

##### rsfMRI data preprocessing

Preprocessing of the rsfMRI data was performed in MATLAB R2016a (RRID:SCR_001622) using DPARSF (version 4.4_180801, RRID:SCR_002372), which is part of DPABI (version 4.0_190305, RRID:SCR_010501) and uses functions of SPM12 (version 6906, RRID:SCR_007037). Preprocessing included the following steps: (1) slice time correction using the middle slice as a reference, (2) realignment using a six-parameter (three translations and three rotations) rigid body transformation, (3) coregistration of rsfMRI data and the T1-weighted anatomical image, (4) segmentation of the T1-weighted anatomical image into gray matter, white matter, and cerebrospinal fluid (CSF), and estimation of deformation field for spatial normalization, (5) general linear model-based removal of nuisance covariates including (i) low-frequency trends (first degree polynomial), (ii) effects of head motion estimated by the six realignment parameters and their first temporal derivatives, (iii) five principle components of white matter and cerebrospinal fluid signals using CompCor (Behzadi et al., 2007), and (iv) the global signal, (6) temporal filtering (0.008–0.09 Hz), (7) spatial normalization of rsfMRI data to MNI space using DARTEL (Ashburner, 2007), (8) interpolation to an isotropic voxel size of 3 mm^3^, (9) spatial smoothing using an 8 mm full-width-at-half-maximum (FWHM) kernel, and (10) removal of scans (“scrubbing”) with framewise displacement (FD) ≥ 0.5 mm, together with the scan immediately before, and together with the two scans immediately after the scan with FD ≥ 0.5 (Power et al., 2012). The quality of spatial normalization was manually inspected.

##### DWI data preprocessing

Preprocessing of the DWI data was performed in FSL (version 6.0.1, RRID:SCR_002823). First, we used *topup* to estimate susceptibility-induced and eddy current-induced distortions based on the non-diffusion weighted images acquired in opposing phase encoding directions. Then, we simultaneously corrected for these distortions and for motion artifacts using *eddy* (Andersson and Sotiropoulos, 2016). As a quality control step, we visually checked the orientation of the principal eigenvector (V1) using *DTIFIT* on the preprocessed DWI data.

##### rsfMRI seed-to-voxel analyses

We examined intra- and interhemispheric functional connectivity between auditory regions of interest (ROIs) and voxels in the temporal, parietal, and frontal lobe. In both hemispheres, the Heschl’s gyrus (HG) and the planum temporale (PT) were selected as seed regions. For each participant, we initially computed the functional connectivity between the seed ROIs and all other voxels of the brain using DPABI. The ROIs were based on probability maps of parcels included in the Harvard-Oxford cortical atlas (probability threshold = 25 %). Functional connectivity maps were built by computing the Pearson correlation coefficient between the preprocessed, spatially averaged time-series within an ROI and the preprocessed time-series of all voxels. To improve the normality of the resulting voxel-wise correlation values, we subsequently applied a Fisher’s r-to-z transformation. This resulted in four (one per ROI) z-transformed connectivity maps per participant, which were subjected to second-level analyses.

##### Group comparisons of functional connectivity maps

To assess the effect of AP, we compared the functional connectivity maps between AP musicians and non-AP musicians. To assess the effect of musicianship, we compared the functional connectivity maps between non-AP musicians and non-musicians. To replicate potential effects of musicianship, we additionally compared AP musicians and non-musicians. For all group comparisons we used nonparametric two-sample *t*-tests (threshold-free cluster enhancement [TFCE] inference, 10,000 permutations) in PALM (version alpha115, RRID:SCR_017029) (Winkler et al., 2014). The significance level was set to α = 0.05, family-wise error (FWE)-adjusted for multiple comparisons. We restricted the search space of the group comparisons using a mask that included the following bilateral regions of the Harvard-Oxford cortical atlas thresholded at 10% probability: HG; PT; planum polare; superior temporal gyrus (STG; anterior and posterior division); middle temporal gyrus (MTG; anterior and posterior division); insular cortex; supramarginal gyrus (SMG; anterior and posterior division); angular gyrus; superior parietal lobule; postcentral gyrus (postCG); precentral gyrus (preCG); inferior frontal gyrus, pars opercularis (IFG,po); inferior frontal gyrus, pars triangularis; middle frontal gyrus; superior frontal gyrus. The selection of these regions was primarily guided by dual-stream models of auditory processing, which, in broad terms, propose that auditory information is processed in two streams: a ventral stream projecting from primary auditory areas on the supratemporal plane along anterior and middle temporal regions to inferior frontal cortex, and a dorsal stream projecting from primary areas along posterior temporal regions to parietal and superior frontal cortices (Rauschecker and Scott, 2009; Leipold et al., 2019b). We also included the insula as its functional connectivity has been previously studied as a function of musicianship (Zamorano et al., 2017; Gujing et al., 2019).

##### Functional connectivity-behavior associations

We used regression analysis for relating behavioral measures of musical aptitude (AMMA total scores), tone-naming proficiency, and musical experience (age of onset of musical training, years of training, cumulative training) to the functional connectivity of the auditory ROIs. Separately for each behavioral measure, we performed voxel-wise regression of the functional connectivity maps with the respective behavioral measure as a single regressor using PALM (TFCE inference, 10,000 permutations, same search space as for the group comparisons). Musical aptitude can be sensibly measured in all participants (Gordon, 1989). However, tone naming requires knowledge on tone names, which non-musicians might not have, and measures of musical experience are only meaningful for musicians. Thus, we included all participants in the voxel-wise regression using the AMMA total scores but only included the musician groups for the regression using the tone-naming scores, age of onset, years of training, and cumulative training. The significance level was set to α = 0.05, FWE-adjusted for multiple comparisons.

##### rsfMRI whole-brain graph-theoretical analysis

To assess effects of AP and musicianship on whole-brain functional connectivity, we used graph theory to characterize global differences in network topology between the groups. For each participant, we computed functional connectivity between all 96 parcels of the Harvard-Oxford cortical atlas (probability threshold = 25 %) using DPABI. Functional connectivity was quantified as Fisher’s r-to-z-transformed Pearson correlation coefficients between the preprocessed, spatially averaged time-series of each parcel. This resulted in a 96 x 96 connectivity matrix per participant representing a whole-brain functional connectome comprising the individual parcels as nodes and the correlation coefficients as edges. Negative edges and edges from the diagonal of the connectivity matrices were set to zero.

Whole-brain functional network topology was quantified using the graph-theoretical measures of *average strength, global efficiency, clustering coefficient, modularity,* and *(average) betweenness centrality* as implemented in the Brain Connectivity Toolbox (version 2019-03-03, RRID:SCR_004841) in MATLAB R2017b (Rubinov and Sporns, 2010). *Average strength* characterizes how strongly the nodes are connected within a network and was defined as the mean of all node strengths. Node strength was computed by taking the sum of all edges of a node. *Global efficiency*, being inversely related to the *characteristic path length*, represents a measure of network integration and was computed as the mean inverse shortest path length in the network. The *clustering coefficient* is a measure of network segregation and was based on *transitivity,* which is the ratio of triangles to triplets in the network. *Modularity* describes the degree to which a network is subdivided into groups of nodes with a large number of within-module edges and a small number of between-module edges. The *(average) betweenness centrality* of the network was defined as the mean nodal betweenness centrality, which itself was computed based on the normalized number of all shortest paths in the network passing through a node.

For each participant, we proportionally thresholded and binarized the connectivity matrices using a wide range of thresholds from 35 % to 1 % retained edges in the network (in steps of 1 %). We then computed the above-listed measures for each threshold resulting in 35 values per measure and participant (*average strength* was based on non-binarized connectivity matrices).

##### Group comparisons of whole-brain functional network topology

Group comparison of the graph-theoretical measures was performed using cluster-based permutation testing in R. Cluster-based permutation testing uses the dependency of graph-theoretical measures across thresholds to control the FWE rate and circumvents the choice of a single arbitrary threshold (Langer et al., 2013; Drakesmith et al., 2015; Brauchli et al., 2020). We estimated the probability of clustered differences between the groups (i.e. across contiguous thresholds) under the null distribution. As before, we separately assessed the effects of AP (by comparing AP to non-AP musicians) and musicianship (by comparing non-AP to non-musicians). In addition, we replicated the potential effects of musicianship by comparing AP to non-musicians. In detail, we first conducted a two-sample Welch’s *t*-test at each threshold. Second, we repeated the first step 5,000 times with permuted group labels. Crucially, we preserved the dependency across thresholds by keeping the random assignment of group labels identical across thresholds within one permutation. Third, we applied a (descriptive) cluster-defining threshold of *p* < 0.05 to build clusters of group differences. Finally, we compared the largest empirical cluster sizes *k* to the null distribution of cluster sizes derived from the permutations. The *p*-value was defined as the proportion of cluster sizes under the null distribution that was larger than or equal to *k* (α = 0.05, FWE-adjusted across multiple thresholds).

##### Whole-brain functional network topology-behavior associations

We assessed associations between the graph-theoretical measures and the behavioral measures (AMMA total scores for all participants; tone-naming proficiency, age of onset, years of training, and cumulative training for the musician groups). For this, we computed the Pearson correlation coefficient (*r*) between the graph-theoretical measure averaged across all thresholds and the particular behavioral measure (α = 0.01, Bonferroni-adjusted across multiple graph-theoretical measures).

##### rsfMRI whole-brain network-based statistic (NBS) analysis

To characterize local between-group differences in the whole-brain functional networks, we identified subnetworks differing between AP and non-AP musicians, between non-AP and non-musicians, and additionally between AP and non-musicians using two-sample *t*-tests as implemented in the network-based statistic (NBS) toolbox (version 1.2, RRID:SCR_002454) (Zalesky et al., 2010). Analogous to cluster-based permutation testing, the NBS approach estimates the probability of group differences in subnetwork sizes under the null distribution and controls the FWE rate on the level of subnetworks. We used the following parameters: 5,000 permutations, test statistic = network extent, and subnetwork-defining thresholds; *t* = 2.8 for AP vs. non-AP, and non-AP vs. non-musicians; and *t* = 3.4 for AP vs. non-musicians. Statistically significant subnetworks were visualized using BrainNet Viewer (version 1.63, RRID:SCR_009446).

##### rsfMRI whole-brain classification analysis

Next, using multivariate pattern analysis (MVPA), we attempted to classify the participants into the three groups based on the individual whole-brain functional connectomes. Group classification of the participants was performed with functions from scikit-learn (version 0.21.2, RRID:SCR_002577) in Python 3.7.0 (RRID:SCR_002577). We first performed a multi-class classification into the three groups (AP, non-AP, non-musicians) using a “one-against-one”-approach with linear support vector machines (C = 1) as classifiers. For each participant, we extracted and flattened the upper right triangle of the connectivity matrix (excluding the diagonal) to build a 4,560-dimensional feature vector representing all edges in the wholebrain functional network. These vectors were associated with their respective group labels (AP, non-AP, non-musician) and stacked to build a dataset. We then z-transformed the dataset per feature and subsequently performed the classification of the participants into the groups. Classification accuracy was estimated using a 5-fold stratified cross-validation. Statistical significance of this accuracy was assessed by repeating the multi-class classification 5,000 times with permuted group labels. The *p*-value was defined as the proportion of accuracies derived from the permutations that were larger than or equal to the empirically obtained accuracy (α = 0.05). To descriptively determine if a small number of features was sufficient for a successful classification, we used recursive feature elimination (RFE), which recursively prunes the least important feature (step = 1) to characterize accuracy as a function of the number of (informative) features (De Martino et al., 2008). The optimal number of features was determined using a 5-fold stratified cross-validation. Subsequently, we performed two follow-up classifications to differentiate AP from non-AP musicians and non-AP from non-musicians. The success of these classifications was quantified by classification accuracy, precision, and recall. We used the identical algorithm, cross-validation scheme, assessment of the statistical significance of the accuracy, and RFE as in the multi-class classification.

#### DWI ROI-to-ROI analysis

Based on the findings from the rsfMRI seed-to-voxel analyses, we next examined the interhemispheric *structural* connectivity between the left and the right PT in the three groups. First, we estimated diffusion parameters based on the preprocessed DWI data by fitting a diffusion tensor model at each voxel using *DTIFIT* in FSL. We specifically focused on two commonly investigated diffusion measures: fractional anisotropy (FA) and mean diffusivity (MD; computed as the mean of the three eigenvalues L1, L2, and L3). Second, we individually reconstructed the white-matter pathways between the left and right PT using probabilistic tractography in FSL (default parameters unless otherwise stated). For this, we fitted a probabilistic diffusion model at each voxel using *BEDPOSTX* (Behrens et al., 2003). Probabilistic tractography was performed on the output of *BEDPOSTX* using *PROBTRACKX* (10,000 samples).

As in the rsfMRI analyses, the ROIs for the probabilistic tractography were based on atlases in MNI space. The seed and target ROIs for the bilateral PT were chosen based on the Harvard-Oxford atlas (probability threshold = 25 %). As a waypoint ROI, we used the midsagittal slice (3 mm thickness) of the corpus callosum map from the Jülich histological atlas (probability threshold = 10 %). As exclusion ROIs, we used the pre- and postcentral gyri as included in the Harvard-Oxford atlas (probability threshold = 25 %) to avoid false-positive pathways terminating in these brain regions. All ROIs were spatially dilated (5 mm spherical kernel) to increase the trackability of the pathways between them and to compensate for interindividual anatomical variability. Because probabilistic tractography was performed in participant-specific diffusion space, we computed the linear transformation from the individual diffusion space to the individual anatomical space using *flirt* and the nonlinear transformation from individual anatomical space to MNI space using *fnirt* in addition to *flirt.* Then, we concatenated these transformations using *convertwarp* and inverted the concatenated transformation using *invwarp*. The resulting warp fields (individual diffusion to MNI space and vice versa) were used in the tractography.

Third, we extracted FA and MD values from the *DTIFIT* output based on the pathways identified by the tractography, more specifically based on the sum of the connectivity distributions of pathways connecting the left PT to the right and vice versa. Before the extraction, we thresholded and binarized the connectivity distributions to retain the 3 % voxels with the highest probability per participant. The extracted FA and MD values were compared between AP and non-AP musicians, and non-AP and non-musicians using Welch’s *t*-tests in R (α = 0.025, Bonferroni-adjusted for multiple diffusion measures). Again, we also compared AP and non-musicians to replicate the potential effects of musicianship. We also associated the FA and MD values with the behavioral measures (AMMA total scores for all participants; tone-naming proficiency, age of onset, years of training, and cumulative training for the musician groups) using *r* (α = 0.025).

#### DWI whole-brain graph-theoretical analysis

Analogously to the rsfMRI analyses, we assessed the effects of AP and musicianship on whole-brain structural connectivity. For this, we performed probabilistic tractography between all parcels of the Harvard-Oxford cortical atlas (probability threshold = 25 %) using *BEDPOSTX*and *PROBTRACKX*(5,000 samples). For each participant, this resulted in a 96 x 96 connectivity matrix representing a whole-brain structural connectome with the parcels as nodes and the connection probability (represented by the number of streamlines) between them as edges. Based on these connectivity matrices, we quantified and compared whole-brain structural network topology between AP and non-AP musicians, non-AP and non-musicians, and additionally between AP and non-musicians. All subsequent analysis steps were identical compared to the rsfMRI whole-brain graph-theoretical analysis (see above for details). We also performed the same correlations between the graph-theoretical measures and the behavioral measures as described above.

#### DWI whole-brain NBS analysis

We repeated the NBS analysis on the structural connectivity matrices to identify structural subnetworks differing between the groups. Apart from the subnetwork-defining threshold (here: *t* = 2.7 for AP vs. non-AP, and non-AP vs. non-musicians, and *t* = 2.8 for AP vs. non-musicians), we used identical parameters as in the rsfMRI analysis (see above for details).

#### DWI whole-brain classification analysis

We also performed the classification analysis based on the whole-brain structural networks. Apart from the different connectivity matrices, all analysis steps and parameters were identical to the rsfMRI whole-brain classification (see above for details).

### General methodological considerations

To comprehensively assess effects of musicianship and AP on functional and structural networks, we used a variety of methods. The acquisition techniques and analytical approaches employed in this study have relative advantages and limitations, which are detailed in the following.

#### Validity and reliability of functional networks derived from rsfMRI

The crucial advantage of rsfMRI is its unique ability to non-invasively resolve functional connections of the human brain at a high spatial resolution. However, the relation between neuronal activity and the blood oxygenation level-dependent (BOLD) signal measured using rsfMRI is indirect and mediated by blood flow, volume, and oxygenation. Electrophysiological oscillations at the neuronal level are correlated with the slow oscillations in BOLD signal (< 0.1 Hz) that are the basis of functional networks. This correlation is not perfect and leaves considerable variance, which can be explained by noise of (non-neuronal) biological and technical origin (Drew et al., 2020). The reliability of functional networks derived from rsfMRI varies greatly depending on factors such as data quantity and quality, brain regions involved, preprocessing choices, time interval between scans, and the spatial level of analysis (local vs. global). While individual edges can exhibit poor reliability (Noble et al., 2019), whole-brain functional networks are remarkably stable and highly sensitive to interindividual differences (Gratton et al., 2018), making them prime targets for comparing groups of different expertise, e.g., musicians and non-musicians.

#### Validity and reliability of structural networks derived from DWI

At present, DWI is the sole method for the non-invasive investigation of the white-matter pathways underlying structural networks of the human brain in vivo. Concerning neuroanatomical validity, it has been shown that the estimation of fiber orientations based on DWI can be reasonably high, although the measurement is indirect because it is based on water diffusion, and estimation accuracy depends on acquisition parameters (spatial resolution, number of directions), and the conformity between complexity of the studied white-matter architecture and the mathematical model to infer this architecture, among others (Jones et al., 2020). Furthermore, tractography algorithms can lack specificity in identifying whitematter tracts (Maier-Hein et al., 2017), but can also lack sensitivity for certain tracts. For example, Westerhausen et al. (2009) did not identify a tract connecting bilateral PT in more than 10% of participants (see below for similar findings in our study). Finally, the neurobiological interpretation of diffusion measures, e.g., FA and MD, is notoriously challenging as there are no straightforward correlates of these measures in white-matter microstructure, and DWI-based tractography cannot provide a quantitative estimate of connection strength but only an estimate of connection probability (Jones et al., 2013). On the upside, the reliability of structural networks based on DWI is relatively high, but also dependent on many factors, e.g., acquisition parameters (Wang et al., 2012), or preprocessing choices (Madhyastha et al., 2014).

#### Merits and shortcomings of ROI-based and whole-brain analysis approaches

Focusing on a set of brain regions in a seed-to-voxel analysis or separate tracts in an ROI-based approach is well-suited to test specific hypotheses and alleviate the multiple-comparisons problem. On the other hand, whole-brain approaches are more suitable for exploration and discovery. Combining both approaches, as we have done in this study, provides a more complete picture than using each approach on its own. The same applies to the use of separate flavors of whole-brain approaches, which in turn have relative advantages and limitations. First, using graph theory has the advantage that the same approach can be applied to both functional and structural networks, providing metrics that quantify topological features of these networks in a single or a few values (Rubinov and Sporns, 2010). Graph-theoretical measures provide a bird’s-eye view of networks that complements the focused perspective of ROI approaches. An issue with graph theory concerns the use of thresholding to remove spurious connections: The type of thresholding employed in graph-theoretical analyses of brain networks (e.g., proportional or absolute thresholding) is subject to ongoing discussions (Van Wijk et al., 2010; van den Heuvel et al., 2017). Absolute thresholding can lead to group differences in the number of edges in the networks which in turn causes spurious group differences in topology (Van Wijk et al., 2010). Proportional thresholding, as used here, equates the number of edges in the network but has been criticized for being sensitive to overall differences in functional connectivity, especially in the presence of potentially random edges (van den Heuvel et al., 2017). Nonetheless, global graph-theoretical measures show high reliability in functional (Termenon et al., 2016) and structural networks (Owen et al., 2013). Second, the application of NBS to whole-brain networks offers the opportunity to identify subnetworks differing between groups without having to test each connection separately. This allows for the localization of connectivity differences that might drive connectivity differences on the global, connectome level. On the downside, NBS is also threshold-dependent and group differences in individual edges should not be interpreted on their own but only in the context of the whole subnetwork (Zalesky et al., 2010). Finally, seed-to-voxel, graph theory, and NBS analyses, as employed here, are (mass)-univariate in nature and thus sensitive for homogeneous increases and decreases in connectivity or network topology in one group relative to another. In contrast, multivariate approaches based on machine learning algorithms show high sensitivity for group differences in *patterns* of connectivity characterized by simultaneous increases and decreases (Haynes, 2015).

### Results

#### Behavioral results

Participant characteristics are given in Table 1. Group comparisons revealed no differences regarding age (*F*(2,150) = 0.59, *p* = 0.55, BF_01_ = 9.30, η^2^_G_ = 0.008), movement during rsfMRI (*F*(2,150) = 0.97, p = 0.38, BF_01_ = 6.75, η^2^G = 0.01), and movement during DWI (*F*(2,150) = 1.44, p = 0.24, BF_01_ = 4.54, η^2^_G_ = 0.02). Both musician groups showed substantially higher musical aptitude than non-musicians as measured by the AMMA total score; AP musicians vs. non-musicians: *t*(85.22) = 8.48, *p* < 0.001, BF_10_ > 100, *d* = 1.69; non-AP musicians vs. non-musicians (*t*(91.17) = 6.54, *p* < 0.001, BF_10_ > 100, *d* = 1.30). There was a trend towards a higher musical aptitude in AP musicians than in non-AP musicians (*t*(99.12) = 1.99, *p* = 0.05, BF_10_ = 1.21, *d* = 0.39), driven by higher AMMA tonal scores in AP musicians (*t*(98.43) = 2.28, *p* = 0.02, BF_10_ = 2.05, *d* = 0.45). The musician groups were comparable in the AMMA rhythm scores (*t*(99.87) = 1.41, *p* = 0.16, BF_01_ = 1.98, *d* = 0.28). With regard to tone-naming proficiency, AP musicians showed substantially higher tone-naming scores than non-AP musicians (*t*(100.95) = 13.68, *p* < 0.001, BF_10_ > 100, *d* = 2.70), and non-AP musicians showed better tone naming than non-musicians (*t*(53.43) = 5.54, *p* < 0.001, BF_10_ > 100, *d* = 1.11). The musician groups did not differ in their age of onset of musical training (*t*(100.96) = −1.00, *p* = 0.32, BF_01_ = 3.08, *d* = 0.20), years of musical training (*/*(100.91) = 1.53, *p* = 0.13, BF_01_ = 1.71, *d* = 0.30), and lifetime cumulative musical training (*t*(96.81) = 1.13, *p* = 0.26, BF_01_ = 2.74, *d* = 0.22).

#### Group differences in functional connectivity of auditory ROIs

To assess the effects of AP and musicianship on the functional connectivity of the auditory ROIs, we compared the functional connectivity maps between AP and non-AP musicians, and between non-AP musicians and non-musicians (the minimal FWE-corrected *p* values per cluster [*p*_FWE_] and cluster sizes [*k*] are given in brackets). Group comparisons between AP musicians and non-AP musicians revealed no statistically significant clusters for any of the four auditory seed ROIs (all *p*_FWE_ > 0.05). Comparisons between non-AP musicians and non-musicians revealed that non-AP musicians showed increased interhemispheric functional connectivity between the left PT (seed ROI) and a cluster in the right PT (*p*_FWE_ = 0.02, *k* = 47; see Figure 1A). A subset of this cluster also survived additional correction across the four ROIs (*p*_FWE-ROI-corr_. = 0.04, *k* = 7). We also identified differences in the symmetric functional connection between the right PT (seed ROI) and two clusters in the left PT (*p*_FWE_ = 0.03, *k* = 51 and *p*_FWE_ = 0.04, *k* = 8). These clusters did not survive additional correction across ROIs (minimum *p*_FWE-ROI-corr_. = 0.08). Details on the clusters are given in Table 2.

**Figure 1.**
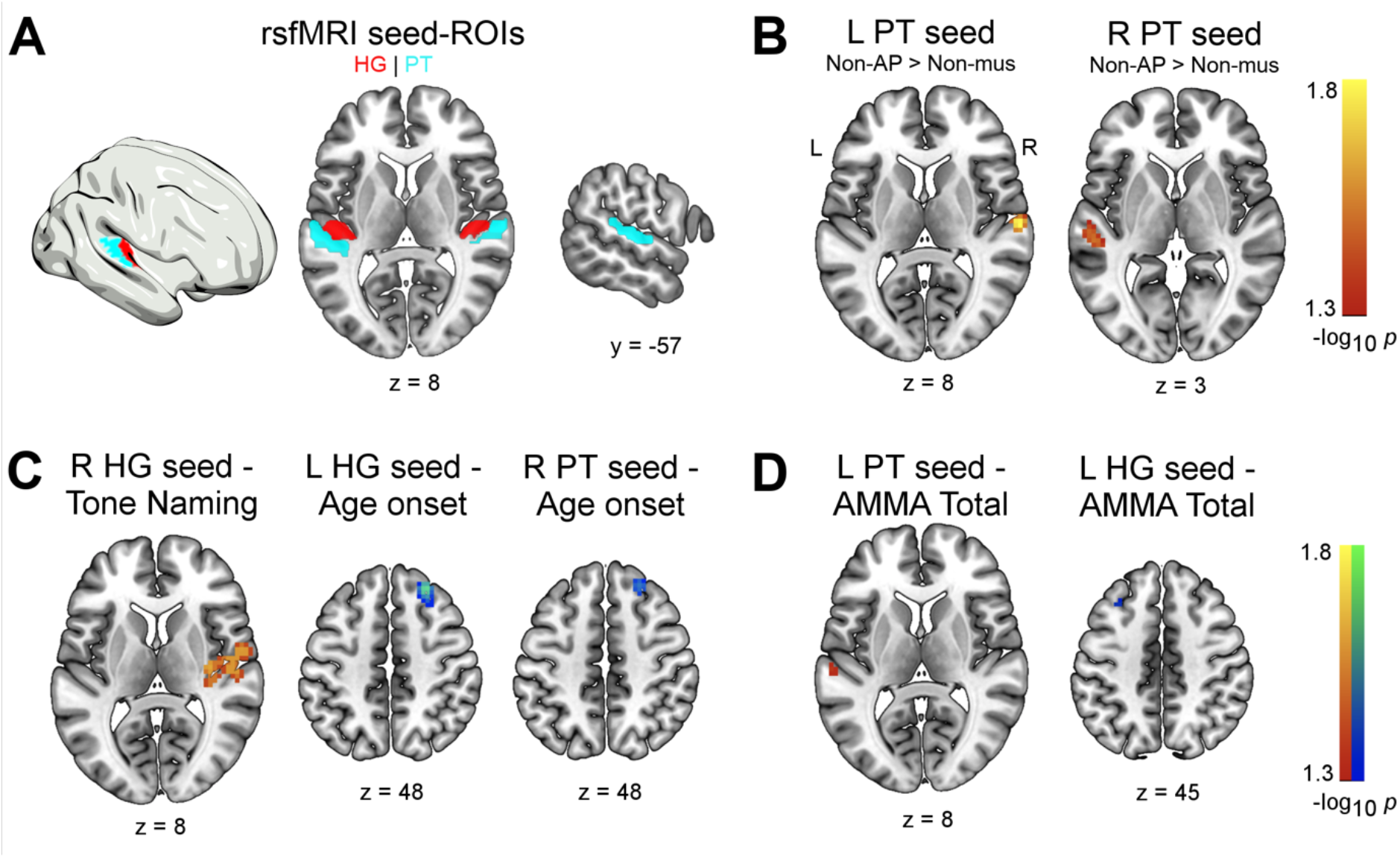
A) Auditory regions of interest (ROIs) used in the rsfMRI seed-to-voxel analyses. Heschl’s gyrus (HG) in red; planum temporale (PT) in sky-blue. Maps for ROIs were derived from the probabilistic Harvard-Oxford cortical atlas as implemented in DPABI. B) Increased intrinsic functional connectivity between left and right PT in non-AP musicians compared to non-musicians (*p*_FWE_ < 0.05). See Extended Data Figure 1-1 for a visualization of group differences in functional connectivity between AP and non-musicians. C) Associations between functional connectivity and behavior in musicians and D) across all subjects (*p*_FWE_ < 0.05). Abbreviations: AMMA = Advanced Measures of Music Audiation; HG = Heschl’s gyrus; L = left; PT = planum temporale; R = right; ROIs = regions of interest.

**Table 2.** Statistically significant group differences between non-absolute pitch musicians and non-musicians in the rsfMRI seed-to-voxel analysis.

| Contrast | Seed region | Target region | $k$ | $p_{\text{FWE}}$ | $x$ | $y$ | $z$ |
| --- | --- | --- | --- | --- | --- | --- | --- |
| Non-AP > Non-mus | Left PT | Right PT | 47 | 0.02 | 63 | -18 | 9 |
| Non-AP > Non-mus | Right PT | Left PT | 51 | 0.03 | -54 | -27 | 3 |
| Non-AP > Non-mus | Right PT | Left PT | 8 | 0.04 | -39 | -36 | 9 |

Abbreviations: Non-AP = non-absolute pitch musicians; Non-mus = non-musicians; $k$ = cluster size in voxels; $p_{\text{FWE}}$ = minimal family-wise error-corrected $p$ value in cluster; PT = planum temporale.

**Table 3.** Significant voxel-wise functional connectivity-behavior associations.

| Behavior | Seed region | Target region | Sign | $k$ | $p_{FWE}$ | x | y | z |
| --- | --- | --- | --- | --- | --- | --- | --- | --- |
| Tone naming | Right Heschl's gyrus | Right posterior insula, auditory association areas | + | 242 | 0.02 | 36 | -15 | 15 |
| AMMA total | Left PT | Left PT | + | 5 | 0.04 | -60 | -24 | 9 |
| AMMA total | Left Heschl's gyrus | Left MTG | - | 6 | 0.04 | -30 | 24 | 48 |
| Age onset | Right Heschl's gyrus | Right DLPFC | - | 46 | 0.02 | 27 | 36 | 48 |
| Age onset | Right PT | Right DLPFC | - | 23 | 0.03 | 24 | 36 | 48 |

Abbreviations: AMMA = Advanced Measures of Music Audiation; DLPFC = dorsolateral prefrontal cortex; $k$ = cluster size in voxels; MTG = middle temporal gyrus; $p_{FWE}$ = minimal family-wise error-corrected $p$ -value in cluster; PT = planum temporale; + = positive association; - = negative association.

As we did not find evidence for group differences between AP and non-AP musicians in the functional connectivity of the auditory ROIs, we attempted to replicate the effects of musicianship that we identified via the comparison of non-AP and non-musicians. For this, we compared the functional connectivity maps between AP musicians and non-musicians. These comparisons revealed that AP musicians also showed increased interhemispheric functional connectivity between the left and right auditory regions (see Extended Data Table 2-1). Overall, these clusters were descriptively larger in number and size, and observable from more seed regions (see Extended Data Figure 1-1).

#### Associations between functional connectivity and behavior

Using voxel-wise regression analysis, we related tone-naming proficiency, musical aptitude, and musical experience to the functional connectivity of the auditory ROIs. Within musicians, higher tone-naming proficiency was associated with increased functional connectivity between the right HG (seed ROI) and surrounding regions including the posterior insula and associative auditory areas (*p*_FWE_ = 0.02, *k* = 242). Most voxels of this cluster also survived additional correction across ROIs (*p*_FWE-ROI-corr_. = 0.03, *k* = 152). Across all participants, we found that higher musical aptitude as measured by the AMMA total scores were associated with increased functional connectivity within the left PT (*p*_FWE_ = 0.04, *k* = 5). Furthermore, we unexpectedly observed that higher musical aptitude was associated with lower functional connectivity between the left HG (seed ROI) and a cluster in the left MTG (*p*_FWE_ = 0.04, *k* = 6). Both of these clusters were very small in size (*k* < 10) and did not survive additional correction across ROIs. Within the musician groups, lower age of onset of musical training was associated with increased functional connectivity between the right HG (seed ROI) and a cluster in the right dorsolateral prefrontal cortex (DLPFC) (*p*_FWE_ = 0.02, *k* = 46). This cluster did not survive additional correction for multiple ROIs. We further found that a lower age of onset was associated with increased functional connectivity between the right planum temporale (seed ROI) and the right DLPFC (*p*_FWE_ = 0.03, *k* = 23). A subset of this cluster just survived additional correction for multiple ROIs (*p*_FWE-ROI-corr_. = 0.046, *k* = 6). Finally, we found no evidence for an association between years of training or cumulative training and the functional connectivity of the auditory ROIs (all *p*_FWE_ > 0.05). Significant associations within musicians are depicted in Figure 1C and across all subjects in Figure 1D.

#### Group differences in functional network topology

Group comparisons of whole-brain functional network topology revealed the following results (FWE-corrected *p* values per cluster [*p*_FWE_] and cluster size across contiguous thresholds [*k*] are given in brackets). We found no evidence for group differences between AP and non-AP musicians in any of the investigated graph-theoretical measures (all *p*_FWE_ > 0.05). However, we observed an effect of musicianship on multiple graph-theoretical measures: We found higher *average strength* (*p*_FWE_ = 0.01, *k* = 35), lower *global efficiency* (*p*_FWE_ = 0.04, *k* = 11), and a higher *clustering coefficient* (*p*_FWE_ = 0.01, *k* = 25) in non-AP musicians than in non-musicians (see Figure 2A). We found no evidence for an effect of musicianship on *modularity,* and *betweenness centrality* of whole-brain functional networks (both *p*_FWE_ > 0.05). Strikingly similar results were obtained by comparing AP and non-musicians, replicating the effects of musicianship on functional network topology (see Extended Data Figure 2-1 for details).

**Figure 2.**
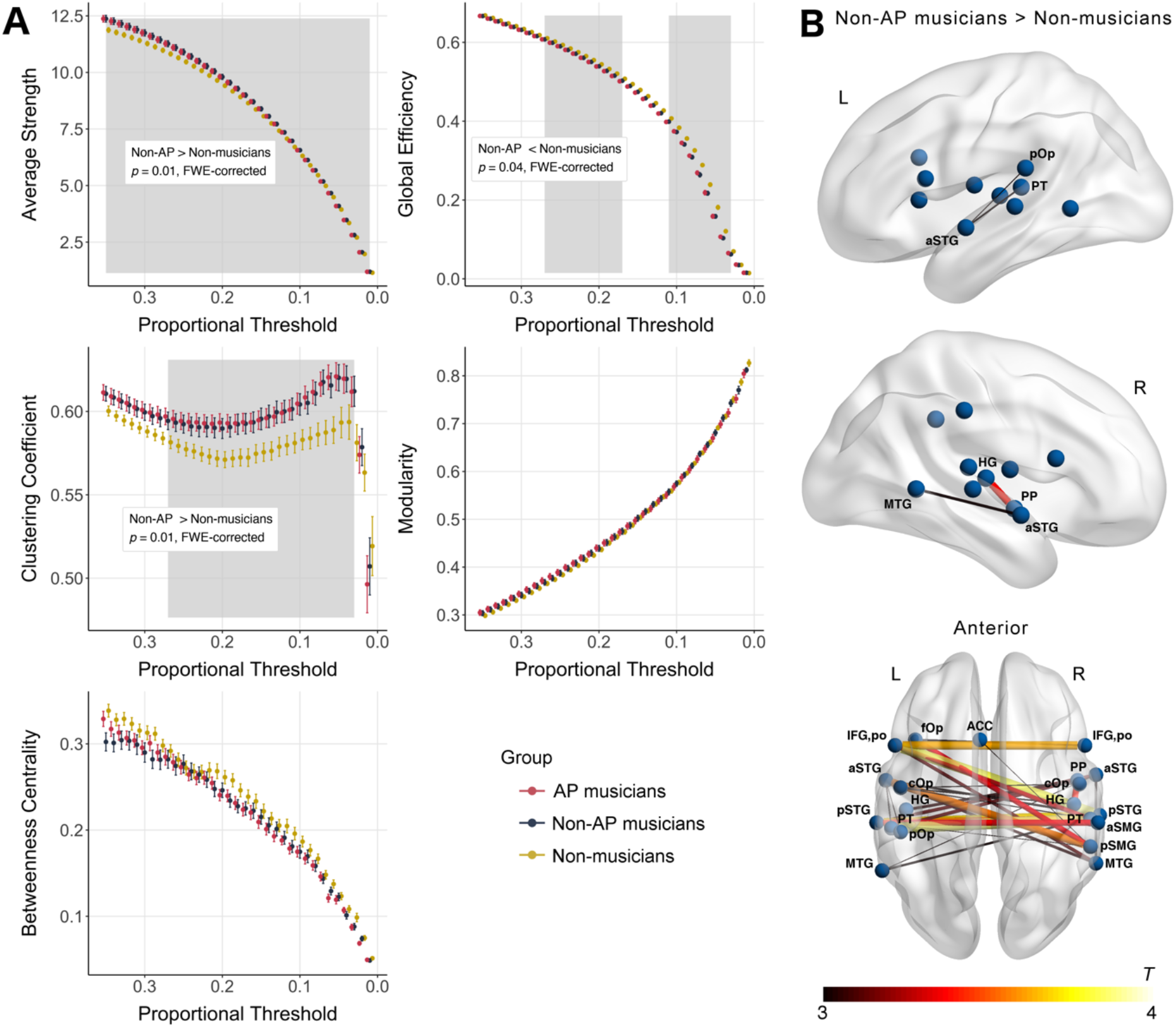
A) Group differences between Non-AP musicians and non-musicians in graph-theoretical measures calculated based on whole-brain functional networks (*p*_FWE_ < 0.05). Gray-shaded area indicates range of thresholds belonging to statistically significant cluster. For details of the comparisons between AP and non-musicians see Extended Data Figure 2-1. B) Subnetwork with increased functional connectivity in non-AP musicians compared to non-musicians obtained in the NBS analysis (*p*_FWE_ < 0.05). Detailed information on all nodes and edges of the functional subnetwork differing between non-AP and non-musicians are given in Extended Data Figure 2-2. For details concerning the functional subnetwork differing between AP and non-musicians see Extended Data Figure 2-3A and Extended Data Figure 2-4. Group classifications based on whole-brain functional networks are visualized in Extended Data Figure 2-5. Abbreviations: ACC = anterior cingulate cortex; AP = absolute pitch; aSMG = anterior supramarginal gyrus; aSTG = anterior superior temporal gyrus; cOp = central operculum; fOp = frontal operculum; HG = Heschl’s gyrus; IFG,po = inferior frontal gyrus, pars opercularis; L = left; MTG = middle temporal gyrus; Non-AP = non-absolute pitch; pSMG = posterior supramarginal gyrus; pSTG = superior temporal gyrus, posterior division; pOp = parietal operculum; PP = planum polare; PT = planum temporale; R = right.

#### Associations between functional network topology and behavior

We found no evidence for an association between *average strength*, *clustering coefficient*, *modularity*, or *betweenness centrality* and any of the behavioral measures for musical aptitude, tone-naming proficiency, or musical experience (all *p* > 0.01 [α = 0.01, adjusted for multiple graph-theoretical measures]). There was a statistically significant negative correlation between *global efficiency* and the AMMA total scores across all participants (*r* = −0.23, *p* = 0.004). However, this correlation was likely driven by group differences in both measures as we found no evidence for a correlation within AP musicians (*r* = 0.01, *p* = 0.90), non-AP musicians (*r* = −0.21, *p* = 0.14), or non-musicians (*r* = −0.11, *p* = 0.49). For all other behavioral measures, we found no evidence for an association with *global efficiency* (all *p* > 0.01).

#### Group differences in whole-brain functional subnetworks

The whole-brain NBS analysis to reveal functional subnetworks differing between the groups did not show evidence for differences between AP and non-AP musicians (*p*_FWE_ > 0.05). In contrast, we identified a subnetwork characterized by higher functional connectivity in non-AP musicians than in non-musicians (*p*_FWE_ = 0.04). As shown in Figure 2B, the descriptively strongest group differences within this subnetwork were present in interhemispheric functional connections between the left and right PT; between the left IFG,po and the right pSTG; between left and right pSTG; and between the left and right IFG,po. Additional nodes of this functional subnetwork were located in brain regions of the temporal and parietal lobes, including HG and anterior and posterior SMG. Detailed information on all nodes and edges of the functional subnetwork differing between non-AP and non-musicians are given in Extended Data Figure 2-2. In the internal replication of these effects of musicianship, we found a strikingly similar subnetwork differing between AP musicians and non-musicians (*p*_FWE_ = 0.005). This functional subnetwork is visualized in Extended Data Figure 2-3A, and details regarding all nodes and edges are given in Extended Data Figure 2-4.

#### Functional network-based classification

Group classification based on whole-brain functional networks using MVPA yielded the following results: The multi-class classification successfully classified the participants into the three groups with an accuracy of 47 %, *p* = 0.002 (chance level = 33 %). See Extended Data Figure 2-5A for a visualization of the null distribution of accuracies with permuted group labels. According to RFE, the optimal number of features for classification was quite large (604 edges), which suggests that the connectivity patterns of a substantial part of the whole-brain functional network contained information about group membership. The confusion matrix showed that the classifier confused AP and non-AP musicians most often, but participants of the musician groups were less often classified as non-musicians and vice versa (see Extended Data Figure 2-5B). Consistent with this pattern of results, the follow-up classification within musicians showed that AP and non-AP musicians could not be successfully differentiated (accuracy = 57 %, *p* = 0.12 [chance level = 50%], precision = 0.56, recall = 0.6; see Extended Data Figure 2-5C). In contrast, the classification of non-AP musicians and non-musicians was successful (accuracy = 65 %, *p* = 0.01 [chance level = 50%], precision = 0.7, recall = 0.6; see Extended Data Figure 2-5D). The optimal number of features necessary for successful classification was again relatively high (1,422 edges).

#### Group differences in transcallosal structural connectivity

In nine AP musicians, 14 non-AP musicians, and 15 non-musicians, probabilistic tractography was not able to identify a white-matter pathway connecting left and right PT (see Figure 3C for a visualization of the white-matter tract). Consequently, these participants were excluded from group comparisons of transcallosal connectivity and the structural connectivity-behavior correlations. Results of the group comparisons of transcallosal structural connectivity are visualized in Figure 3A. We found no evidence for group differences in FA between AP musicians and non-AP musicians (*t*(68.34) = 0.81, *p* = 0.42, *d* = 0.19), and between non-AP musicians and non-musicians (*t*(69.17) = 0.12, *p* = 0.90, *d* = 0.03). Furthermore, there was no evidence for differences in MD between AP and non-AP musicians (*t*(70.02) = −1.01, *p* = 0.31, *d* = 0.23). On the contrary, we found a statistically significant difference in MD between non-AP and nonmusicians, characterized by higher MD values in non-AP than in non-musicians (*t*(59.51) = 2.61, *p* = 0.01, *d* = 0.61). In the internal replication of this effect of musicianship, we found that AP musicians descriptively showed higher MD values than non-musicians, but this difference did not reach statistical significance (*t*(75.11) = 1.81, *p* = 0.07 [α = 0.025, adjusted for multiple diffusion measures], *d* = 0.40).

**Figure 3.**
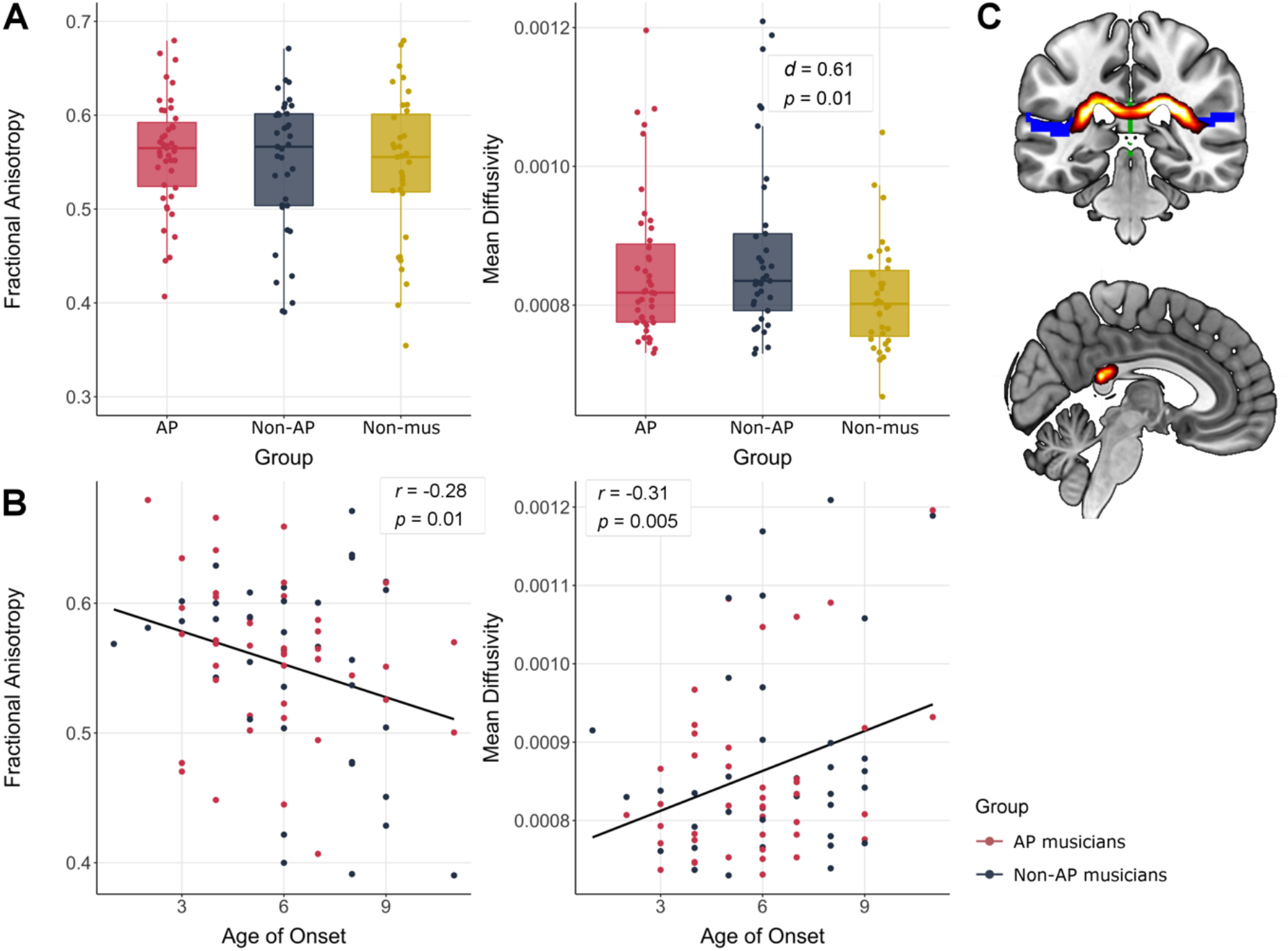
A) Group differences between AP, non-AP, and non-musicians in fractional anisotropy and mean diffusivity values (α = 0.025, adjusted for multiple diffusion measures). B) Associations between fractional anisotropy and mean diffusivity values and age of onset of musical training. C) Coronal and sagittal view of the mean white-matter pathway between left and right planum temporale obtained by probabilistic tractography across all subjects. Abbreviations: AP = absolute pitch; non-AP = non-absolute pitch; Non-mus = non-musicians.

#### Associations between transcallosal structural connectivity and behavior

Structural connectivity-behavior associations are shown in Figure 3B. Across both musician groups, we found a statistically significant negative correlation between the age of onset of musical training and FA values within the pathway connecting left and right PT (*r* = −0.28, *p* = 0.01). We did not find evidence for an association between any of the other behavioral measures and FA (all *p* > 0.025). Furthermore, we found a statistically significant positive correlation between age of onset and MD values across both musician groups (*r* = 0.31, *p* = 0.005). Again, there was no evidence for an association of any of the other behavioral measures and MD (all *p* > 0.025).

#### Group differences in structural network topology

In the analysis of whole-brain structural network topology, we found no evidence for group differences between AP musicians and non-AP musicians, or between both musician groups and non-musicians in any of the investigated graph-theoretical measures (all *p*_FWE_ > 0.05).

#### Associations between structural network topology and behavior

We found a statistically significant positive correlation between *betweenness centrality* and the musicians’ age of onset of musical training (*r* = 0.27, *p* = 0.006). Furthermore, age of onset was also descriptively associated with *mean strength (r* = −0.19, *p* = 0.049), *global efficiency (r* = −0.21, *p* = 0.04), and *clustering coefficient (r* = 0.22, *p* = 0.02; see Figure 4A). However, these correlations did not survive the adjustment of the significance level for multiple graph-theoretical measures. We found no evidence for an association of *modularity* and age of onset. Furthermore, there was no evidence for an association between any of the other behavioral measures (besides age of onset) and the graph-theoretical measures.

**Figure 4.**
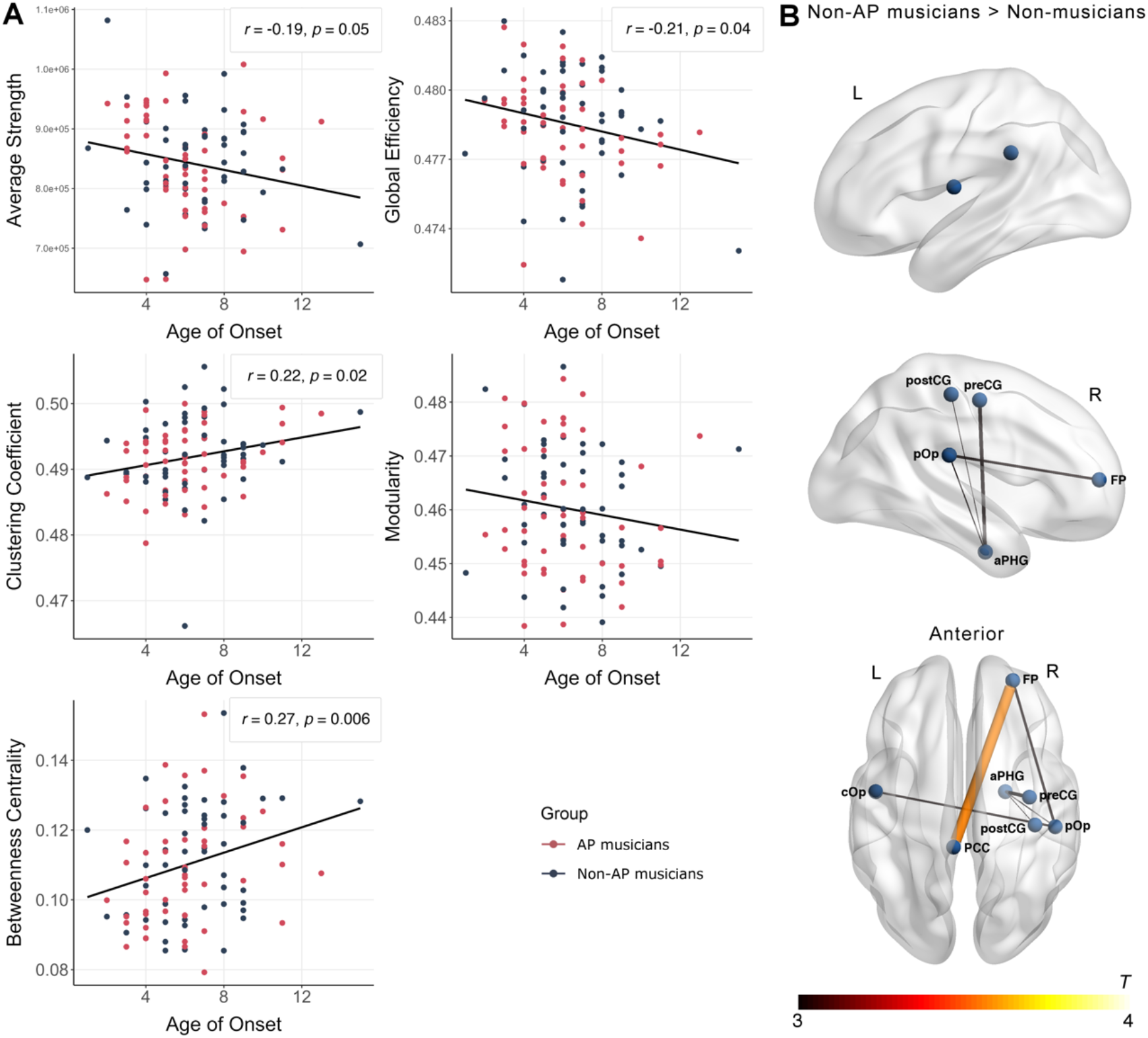
A) Associations between structural network topology and age of onset of musical training for AP- and non-AP musicians. B) Subnetwork with increased structural connectivity in non-AP musicians compared to non-musicians obtained in the NBS analysis (*p*_FWE_ < 0.05). Detailed information on all nodes and edges of the structural subnetwork differing between non-AP and non-musicians are given in Extended Data Figure 4-1. For details concerning the structural subnetwork differing between AP and non-musicians see Extended Data Figure 2-3B and Extended Data Figure 4-2. Abbreviations: AP = absolute pitch; aPHG = anterior parahippocampal gyrus; cOp = central operculum; FP = frontal pole; L = left; Non-AP = non-absolute pitch; PCC = posterior cingulate cortex; postCG = postcentral gyrus; preCG = precentral gyrus; pOp = parietal operculum; PT = planum temporale; R = right.

#### Group differences in whole-brain structural subnetworks

As for the functional data, the NBS analysis to identify structural subnetworks differing between the groups did not show evidence for differences between AP musicians and non-AP musicians (*p*_FWE_ > 0.05). On the contrary, we again identified a subnetwork characterized by higher structural connectivity in non-AP than in non-musicians (*p*_FWE_ = 0.047). As can be seen from Figure 4B, the descriptively biggest group difference in structural connectivity was between the posterior cingulate cortex (PCC) and the frontal pole (FP). Furthermore, non-AP musicians showed higher structural connectivity between right perisylvian regions including the parietal operculum (pOp) as well as preCG and postCG. Detailed information on all nodes and edges of the structural subnetwork differing between non-AP and non-musicians are given in Extended Data Figure 4-1. A similar subnetwork was identified by comparing AP and non-musicians (*p*_FWE_ = 0.003). This subnetwork had descriptively stronger group differences and was more extended than the subnetwork identified by comparing non-AP and non-musicians. This structural subnetwork is visualized in Extended Data Figure 2-3B, and details regarding all nodes and edges are given in Extended Data Figure 4-2.

#### Structural network-based classification

Group classification based on whole-brain structural networks using MVPA yielded no successful classifications. The three groups could not be successfully differentiated in the multi-class classification (accuracy = 35 %, *p* = 0.33 [chance level = 33 %]). Furthermore, the follow-up classifications showed that neither non-AP and AP musicians (accuracy = 43 %, *p* = 0.90 [chance level = 50%], precision = 0.41, recall = 0.49), nor non-AP and non-musicians (accuracy = 52 %, *p* = 0.35 [chance level = 50%], precision = 0.53, recall = 0.52) could be successfully differentiated.

### Discussion

In this study, we assessed the effects of musicianship and AP on brain networks. Our main results are summarized in Table 4. We found robust effects of musicianship across various methodological approaches, which were largely replicable in AP and non-AP musicians. Both musician groups showed stronger interhemispheric functional connectivity between left and right PT, enhanced connectivity in temporal-parietal-frontal functional subnetworks, and globally altered functional network topology, compared to non-musicians. Furthermore, non-AP musicians and non-musicians could be successfully classified using MVPA based on functional connectomes. Musicians also showed altered transcallosal structural connectivity in the white-matter tract connecting bilateral PT. We detected several brain-behavior associations between connectivity and behavioral measures of musicianship, most prominently between structural network features and the age of onset of musical training. Finally, we found no evidence for group differences between non-AP and AP musicians across all analyses: the two musician groups showed striking similarities in both functional and structural networks.

**Table 4.**
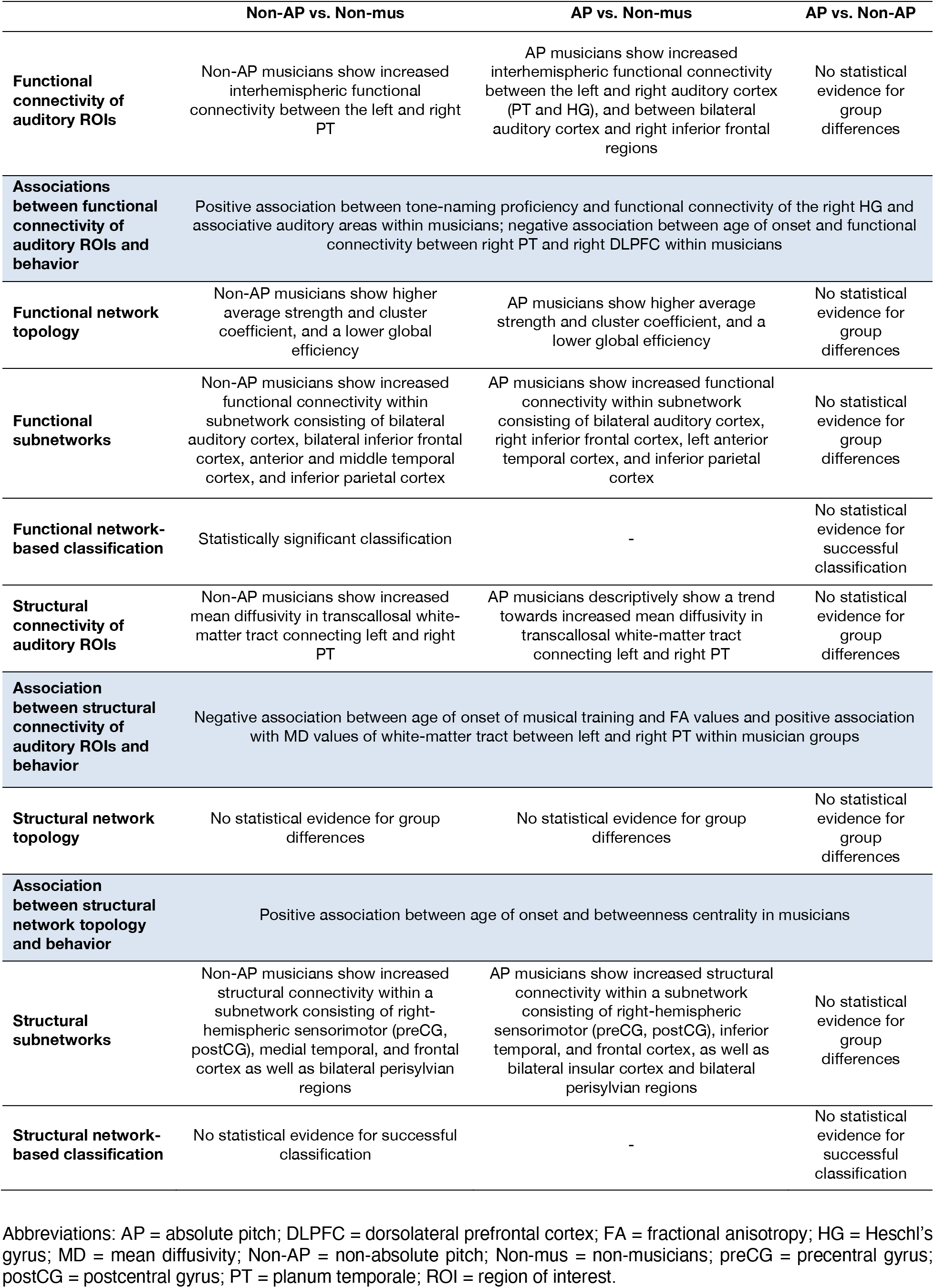
Summary of main findings for group comparisons, classifications, and brain-behavior associations.

Results showed altered connectivity between left and right PT in both musician groups compared to nonmusicians. Left and right PT are structurally connected via the isthmus and splenium of the corpus callosum (Hofer and Frahm, 2006). Whereas effects of musicianship on (more anterior) parts of the corpus callosum have been frequently observed (Schlaug et al., 1995; Bengtsson et al., 2005; Vollmann et al., 2014), only one previous study has reported microstructural differences between musicians and non-musicians in the callosal fibers connecting bilateral PT (Elmer et al., 2016). Here, we showed that altered microstructural connectivity is accompanied by increased intrinsic functional connectivity in musicians, an observation that substantiates earlier reports of increased functional connectivity between bilateral auditory areas using electroencephalography (Klein et al., 2016). The PT’s role in auditory processing is well documented (Griffiths and Warren, 2002). Increased interhemispheric functional connectivity in musicians might reflect increased information transfer between the homotopic areas. It is conceivable that enhanced auditory information coordination is the basis for the superior auditory skills frequently noted in musically trained individuals (Schneider et al., 2002; Kraus and Chandrasekaran, 2010).

The effects of musicianship on functional networks were not restricted to interhemispheric auditory-to-auditory connections: We identified widespread subnetworks showing enhanced connectivity in musicians, mostly encompassing bilateral superior and middle temporal, inferior frontal, and inferior parietal regions. These regions can be well situated within the frameworks of dual-stream models for auditory processing (Rauschecker and Scott, 2009). In particular, our data suggests that communication between regions of the bilateral ventral stream is shaped by musicianship more strongly than that between regions of the dorsal stream (see Figure 2). However, most altered connections in the subnetwork were of interhemispheric nature. It has been shown that interhemispheric information transfer causally modulates expansive auditory and motor networks during rest (Andoh et al., 2015). Thus, experience-dependent plasticity in interhemispheric connections could have a prime role in modulating network interactions between auditory areas and cortical regions in the temporal, parietal, and frontal lobes. As we were able to replicate virtually the same enhanced subnetworks in both non-AP and AP musicians compared to non-musicians, the identified subnetworks of the current study seem to robustly reflect general characteristics of musical expertise.

A notable feature of the DWI results is the consistent and highly specific association between the age of onset of music training and structural network measures. Importantly, these network measures were not associated with other behavioral measures such as cumulative training hours and years of training. Age of onset of musical training was correlated with diffusion measures in the transcallosal white-matter tract connecting left and right PT. This result complements previous reports of associations between age of onset and diffusion measures in parts of the corpus callosum connecting bilateral sensorimotor brain regions (Steele et al., 2013). An earlier study also showed an association of age of onset with diffusion measures of both the anterior and the posterior part of the corpus callosum (Imfeld et al., 2009). These findings suggest that microstructural properties of the corpus callosum are sensitive for changes when musical training starts at a young age, possibly during a sensitive period when the potential for plasticity is especially high (Schlaug et al., 1995). Additionally, for the first time, we observed associations between age of onset and whole-brain structural network topology. Thus, musical training during early childhood not only has local effects on microstructure, but also has global effects on the topology of the structural connectome, and these effects are stronger the earlier musical training begins.

This is the first study to analyze effects of musicianship on both structural and functional connectivity. In this context, we found a surprisingly low correspondence between effects on functional versus structural networks. Evidence suggests that rsfMRI-based functional connectivity and DWI-based structural connectivity are, to some extent, related (Hermundstad et al., 2013). However, because of indirect structural connections, functional connectivity between regions can also be observed without direct structural links (Honey et al., 2009). We found that effects of musicianship on connectivity were particularly strong in the functional domain, and less so in the structural domain. Therefore, based on our data, one might speculate that musical training more strongly shapes functional networks, and does so mostly independently of structural networks. An important exception to this general hypothesis concerns the observed differences in transcallosal connectivity between bilateral PT. However, this selective correspondence is highly consistent with the finding that interhemispheric functional connectivity causally depends on structural connectivity provided via the corpus callosum (Jäncke and Steinmetz, 1994, 1998; Roland et al., 2017).

Concerning reproducibility, the effects of musicianship were not as widespread as one might have expected from previous evidence on brain function and structure in musicians (e.g., Schlaug, 2015). This divergence could be attributable to a number of reasons: First, some previously reported findings might not be reproducible because of inadequate sample sizes (Button et al., 2013). Second, as outlined above (see General methodological considerations), the methodology applied in this and previous studies may lack the reliability for the effects to be consistently observed in different studies. Also, stereotactic normalization might diminish group differences in anatomy (e.g., asymmetries), which could have downstream consequences on connectivity. Future studies will benefit from approaches that consider interindividual anatomical variance (Dalboni da Rocha et al., 2020). Third, the investigation of intrinsic functional and structural networks could be less sensitive compared to activation or connectivity in task-based experiments, e.g., during auditory or motor tasks (Bangert et al., 2006). One possibility to disentangle these potential causes are well-powered replication studies in a collaborative setting, making data acquisition from large samples of musicians feasible. Future studies could also benefit from a hypothesis-driven framework, where brain regions and tracts putatively involved in music production, e.g., the hand area in motor cortex or the arcuate fasciculus, are investigated more closely (Halwani et al., 2011; Rüber et al., 2015).

Across analyses, we found remarkable similarity of networks for the two musician groups, which seems surprising, given that previous studies have reported effects of AP on connectivity. There are multiple reasons potentially contributing to this discrepancy. First, previous evidence for the effects of AP on connectivity is sparse: the number of studies reporting differences in intrinsic functional and structural connectivity is relatively small, none of the effects have been replicated to date, and the effects reported were very subtle in size (Greber et al., 2020). Second, most of the studies investigated small to very small samples, making them prone to false-positive results (Button et al., 2013). Third, methodology varied widely, both between previous studies and compared to the current study. As outlined above (see General methodological considerations), current methodology might lack the sensitivity and reliability to robustly detect subtle differences. Fourth, there is no agreement on defining AP; it might represent a distinct population (Athos et al., 2007) or lie on the upper end of a continuum of tone-naming abilities (Bermudez and Zatorre, 2009). We defined AP based on self-report, and the tone-naming proficiency of our AP and non-AP musicians strongly differed (*d* > 2). Thus, we are confident that the similarities of AP and non-AP musicians are valid. It is important to note that our results should not be regarded as evidence that there are no effects of AP on the brain in general. For example, we found a correlation between tone naming and functional connectivity of right HG and surrounding areas. This is consistent with previous reports of AP-specific alterations in right-hemispheric auditory regions (Leipold et al., 2019a), and underlines the importance of right-hemispheric HG in AP (Wengenroth et al., 2014). Furthermore, task-based studies investigating tone labeling in action have shown considerable promise for uncovering the neural peculiarities of the AP phenomenon (Schulze et al., 2013; Greber et al., 2018; Leipold et al., 2019d, 2019c; McKetton et al., 2019).

To conclude, we identified robust and replicable effects of musical expertise on intrinsic functional and structural brain networks. As effects were stronger in the functional domain, we hypothesize that musical training particularly affects functional compared to structural networks. The effects of AP on large-scale brain networks might be subtle, requiring very large samples or task-based experiments to be detected.

## Acknowledgments

This work was supported by the Swiss National Science Foundation (SNSF), grant no. 320030_163149 to Lutz Jäncke. We are extremely grateful to Désirée Yamada for her invaluable help in data acquisition and research administration. We also thank Silvano Sele for his support with the statistical analyses, Marielle Greber and Lisa-Katrin Kaufmann for helpful comments on the manuscript, and Chantal Oderbolz for proofreading the manuscript.

---

**Extended Data Figure 1-1.**
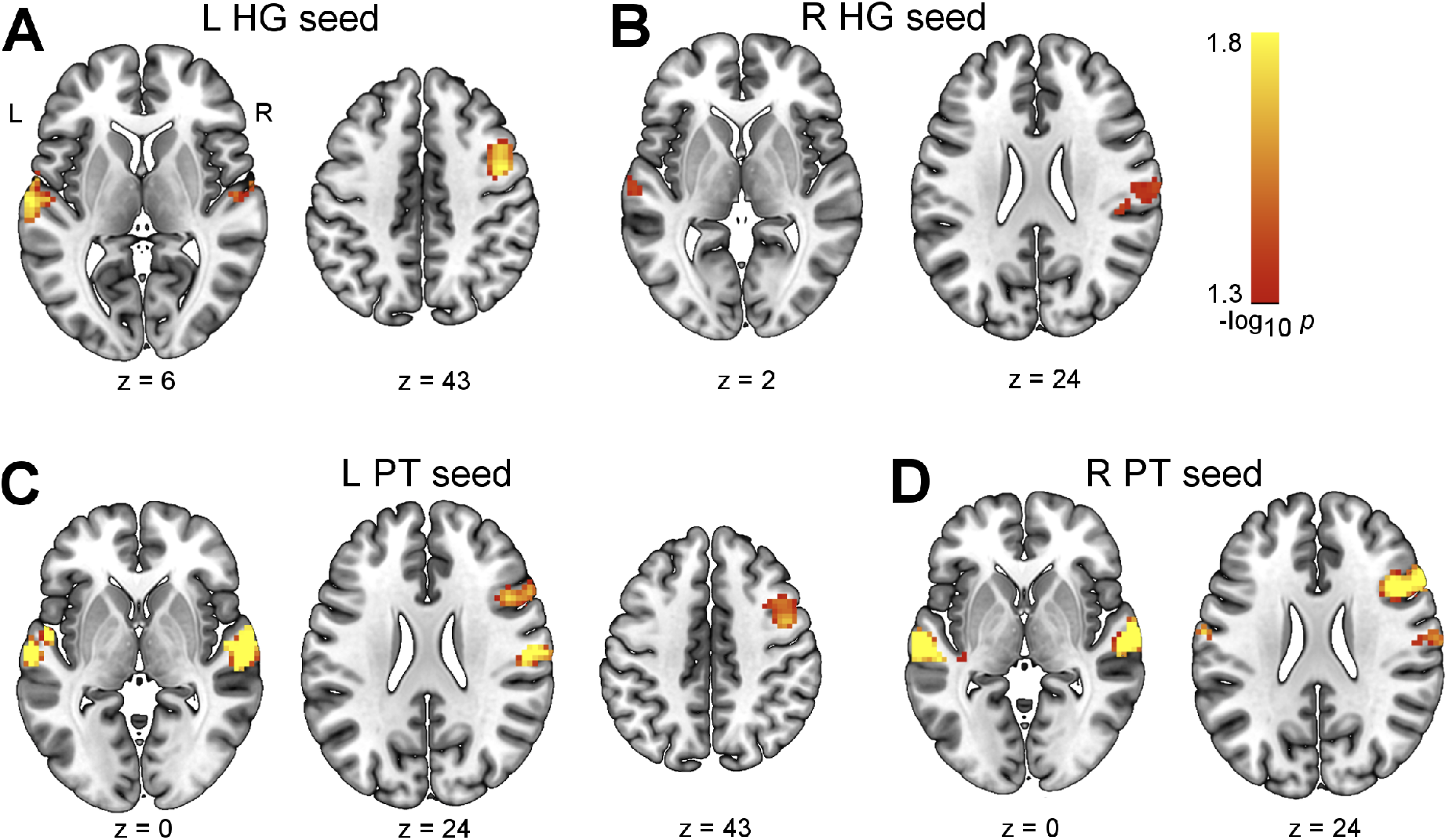
Increased resting-state functional connectivity in **absolute pitch musicians compared to non-musicians** (*p*_FWE_ < 0.05) in the seed-to-voxel analysis for the following seeds: A) the left Heschl’s gyrus, B) the right Heschl’s gyrus, C) the left planum temporale and D) the right planum temporale. Abbreviations: HG = Heschl’s gyrus; L = left; PT = planum temporale; R = right.

**Extended Data Figure 2-1.**
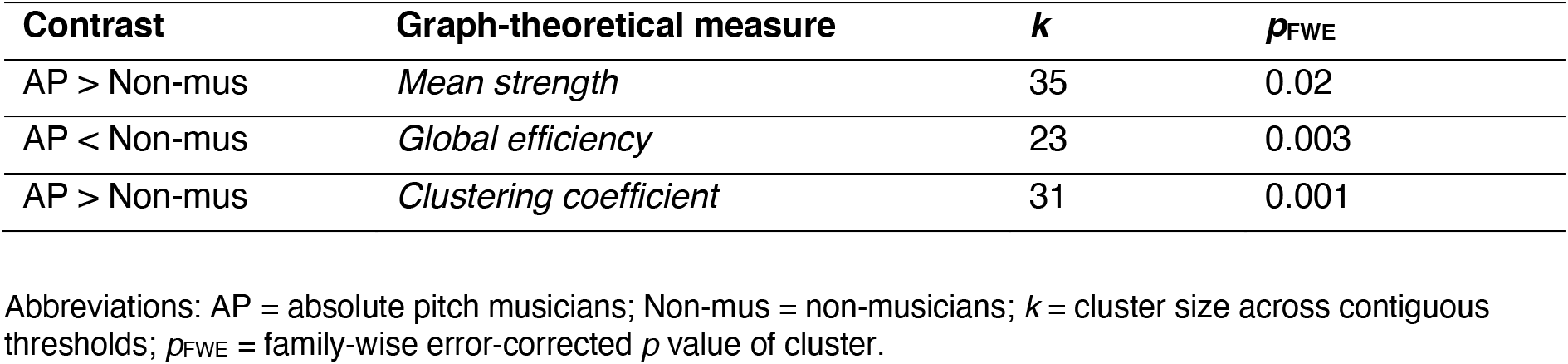
Statistically significant group differences between absolute pitch musicians and non-musicians in whole-brain functional network topology.

**Extended Data Figure 2-2.**
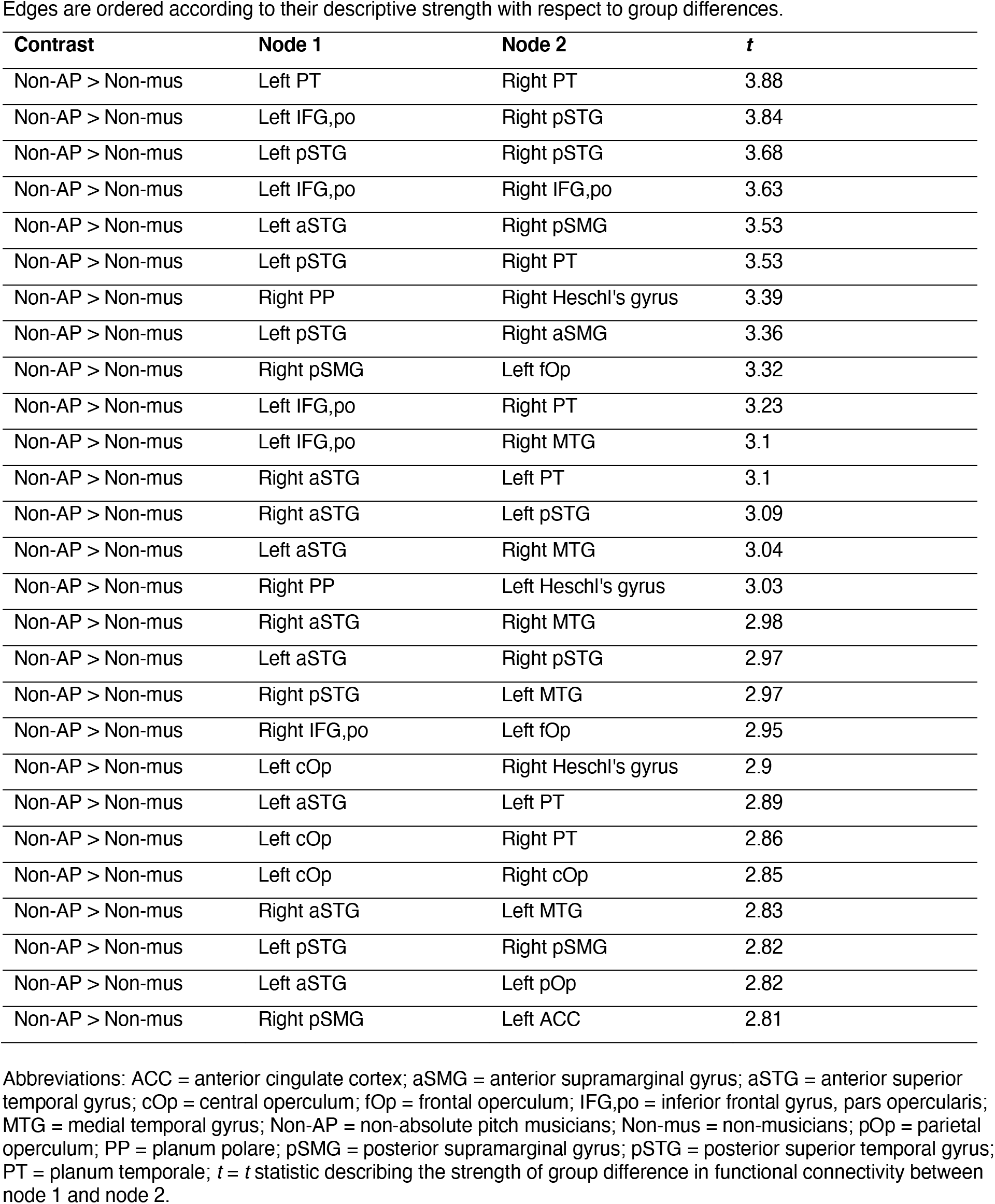
Edges of statistically significant functional subnetwork differing between non-absolute pitch musicians and non-musicians.

**Extended Data Figure 2-3.**
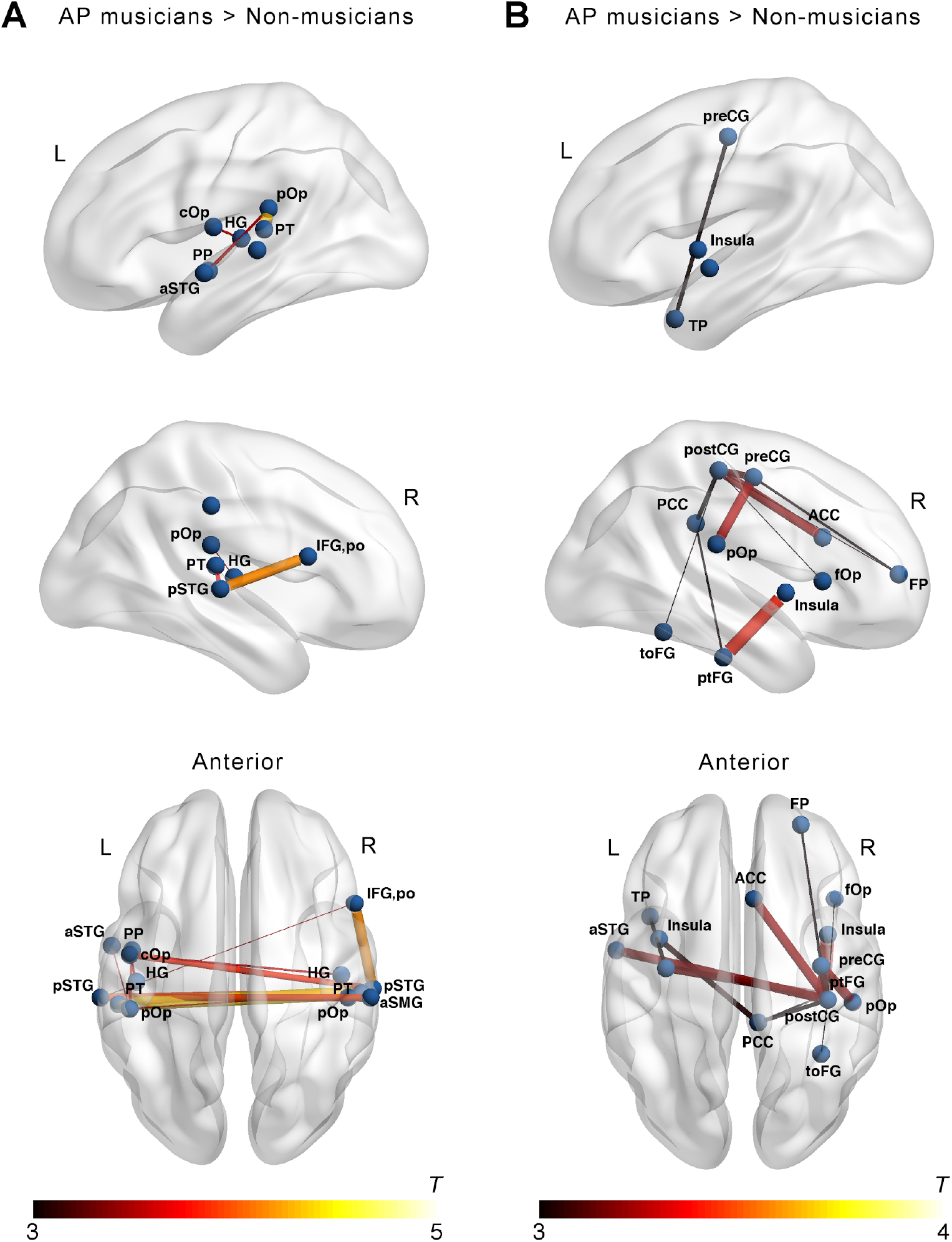
Increased subnetworks in **absolute pitch musicians compared to non-musicians** obtained in the whole-brain network-based statistic (NBS) analysis for A) resting-state functional connectivity and B) diffusion weighted imaging (DWI)-based structural connectivity (*p*_FWE_ < 0.05). Abbreviations: ACC = anterior cingulate cortex; AP = absolute pitch; aSMG = anterior supramarginal gyrus; aSTG = anterior superior temporal gyrus; cOp = central operculum; fOp = frontal operculum; FP = frontal pole; HG = Heschl’s gyrus; IFG,po = inferior frontal gyrus, pars opercularis; L = left; MTG = middle temporal gyrus; PCC = posterior cingulate cortex; postCG = postcentral gyrus; preCG = precentral gyrus; pSTG = superior temporal gyrus, posterior division; pOp = parietal operculum; PP = planum polare; PT = planum temporale; ptFG = posterior temporal fusiform gyrus; R = right; toFG = temporal occipital fusiform gyrus; TP = temporal pole.

**Extended Data Figure 2-4.**
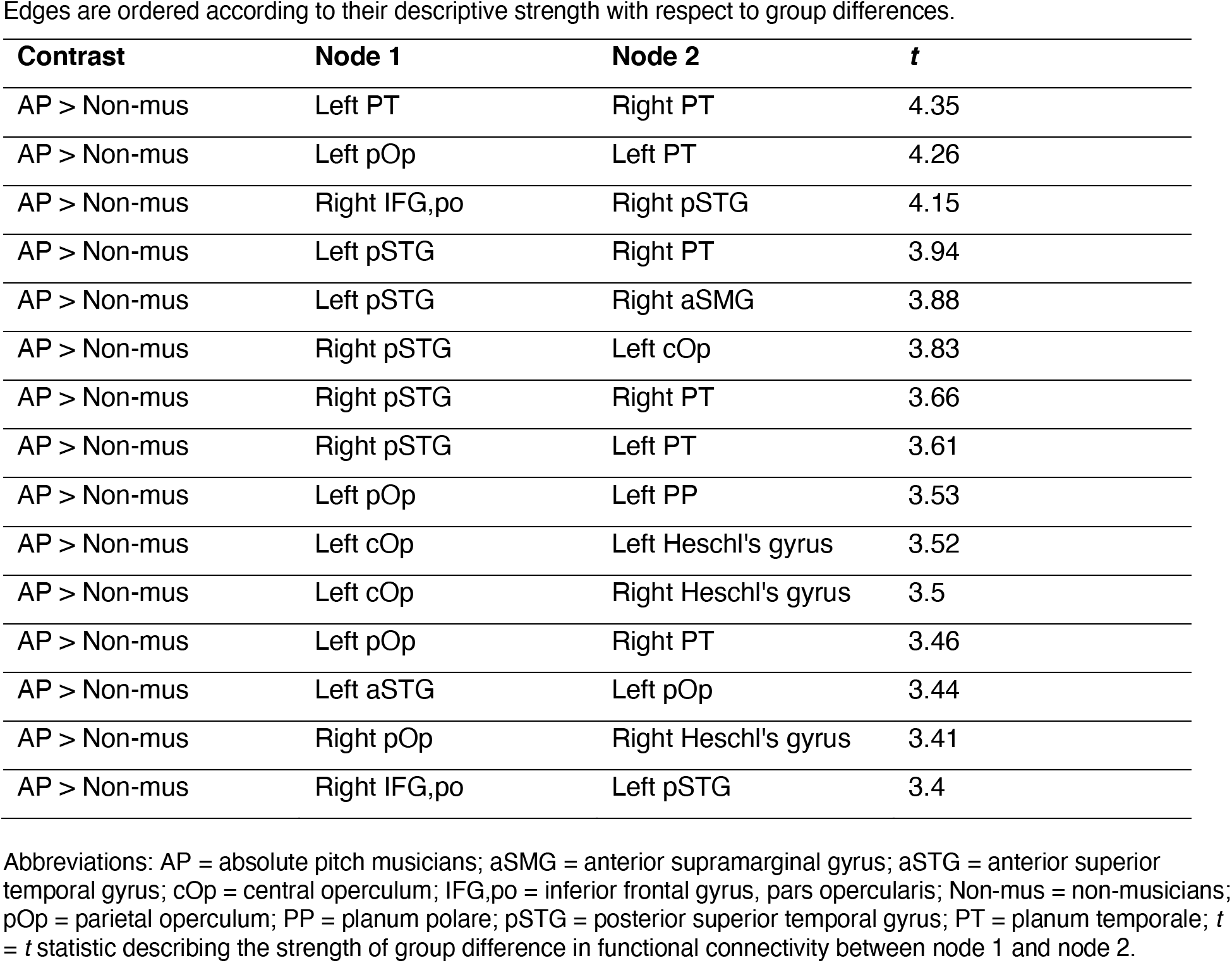
Edges of statistically significant functional subnetwork differing between absolute pitch musicians and non-musicians.

**Extended Data Figure 2-5.**
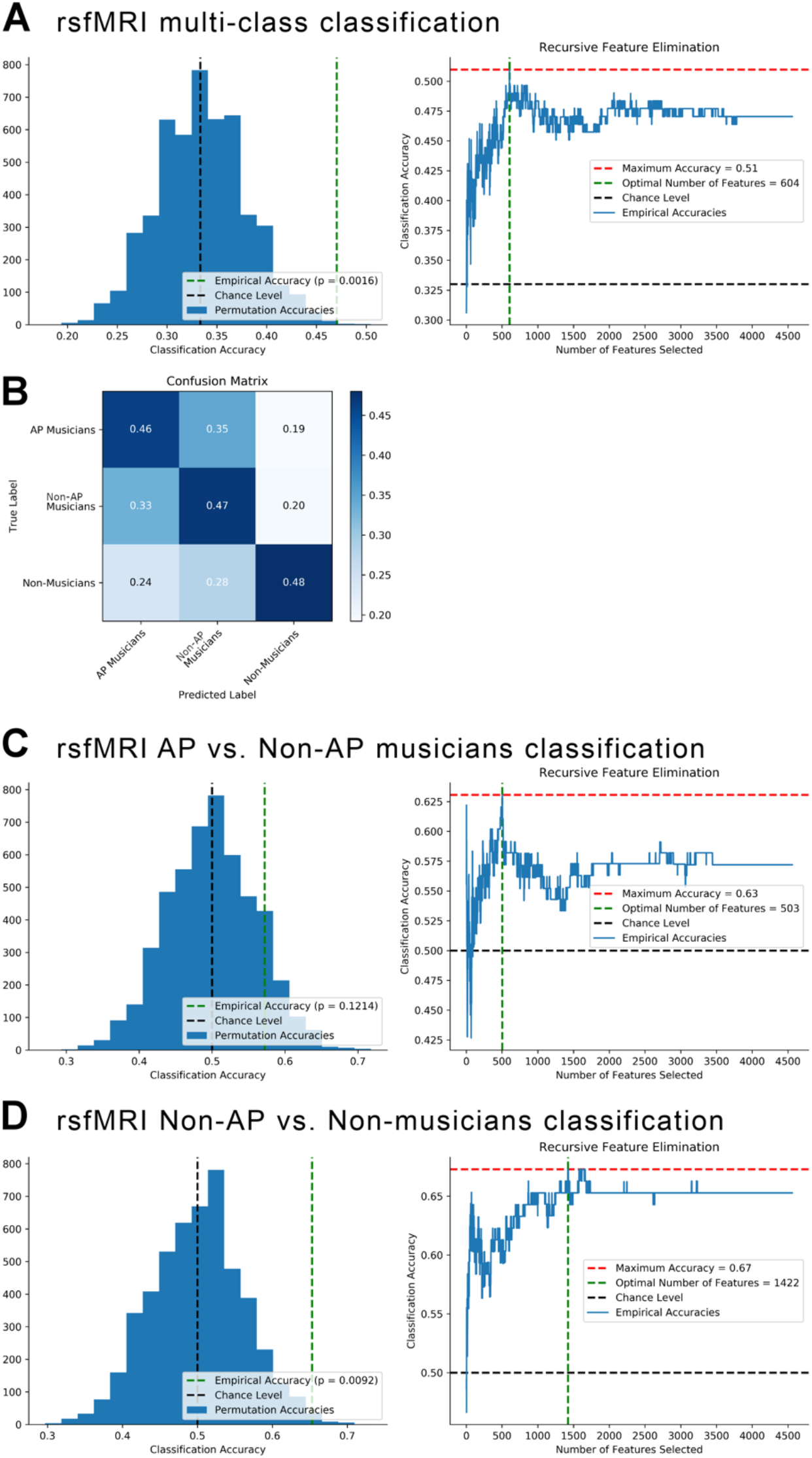
A) Multi-class classification differentiating absolute pitch (AP), non-absolute pitch (non-AP), and non-musicians based on whole-brain functional networks: Null distribution of accuracies with permuted group labels (left) and recursive feature elimination outcome (right). B) Confusion matrix of classifier performance (accuracy) for multi-class classification. C) Classification of AP vs. non-AP musicians: Null distribution of accuracies with permuted group labels (left) and recursive feature elimination outcome (right). D) Classification of non-AP vs. non-musicians: Null distribution of accuracies with permuted group labels (left) and recursive feature elimination outcome (right).

**Extended Data Figure 4-1.**
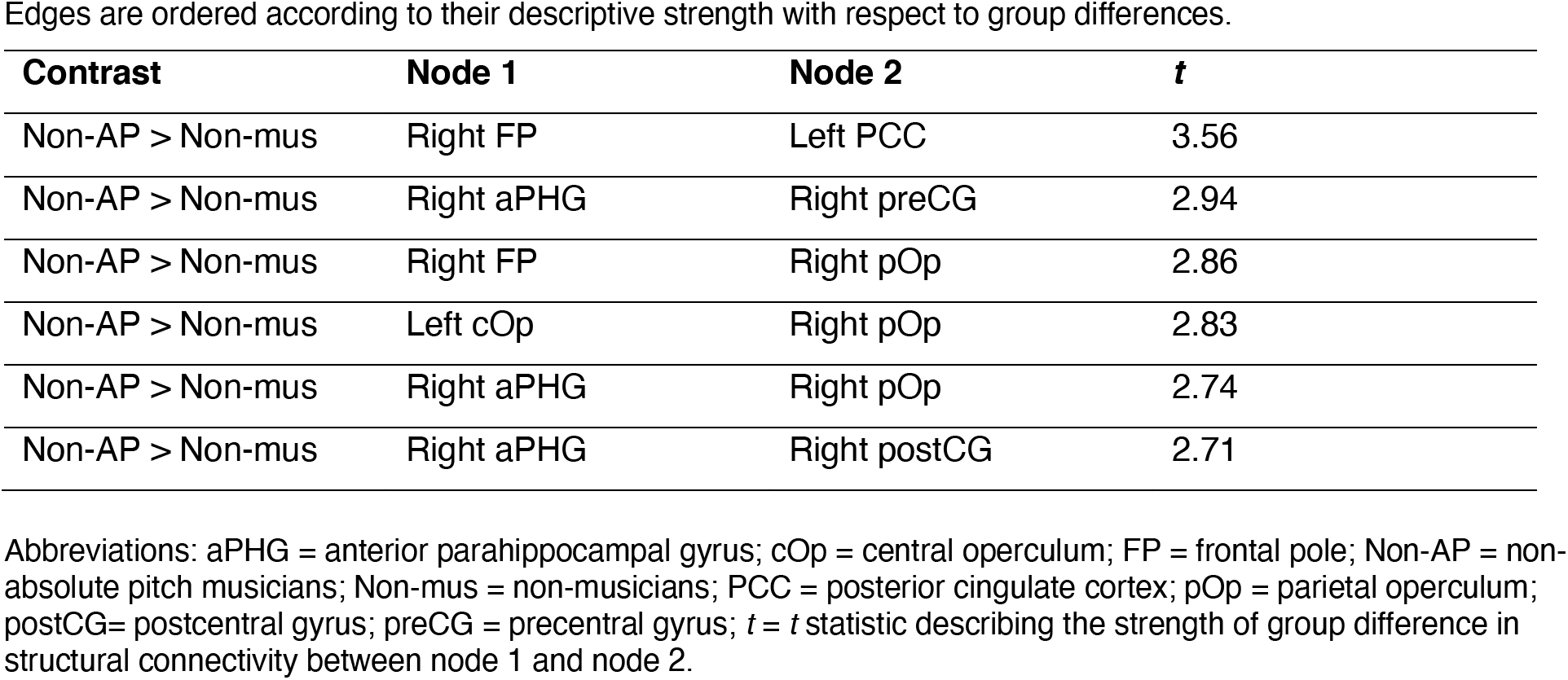
Edges of statistically significant structural subnetwork differing between non-absolute pitch musicians and non-musicians.

**Extended Data Figure 4-2.**
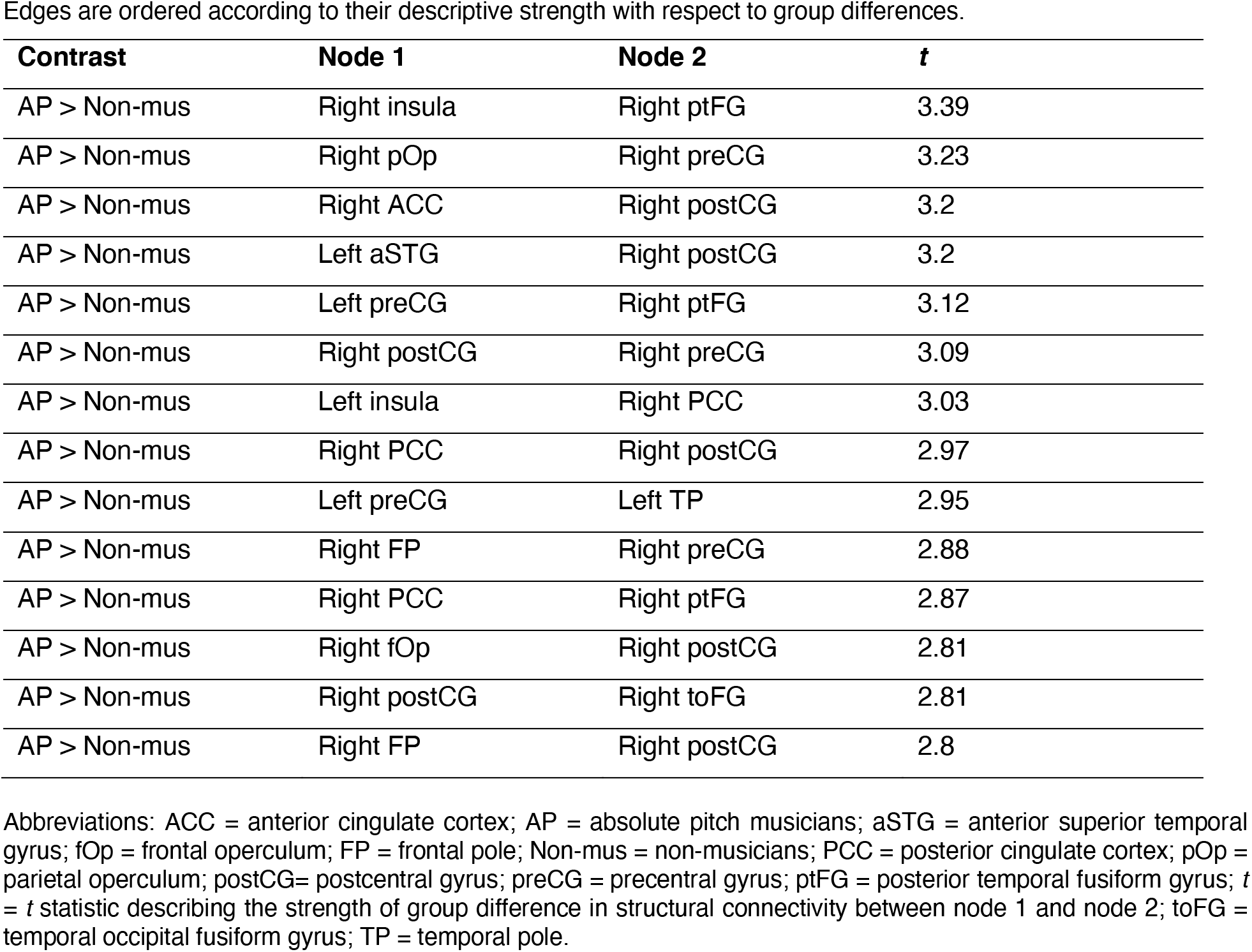
Edges of statistically significant structural subnetwork differing between absolute pitch musicians and non-musicians.

**Extended Data Table 2-1.** Statistically significant group differences between absolute pitch musicians and non-musicians in the rsfMRI seed-to-voxel analysis.

| Contrast | Seed region | Target region | $k$ | $p_{FWE}$ | x | y | z |
| --- | --- | --- | --- | --- | --- | --- | --- |
| AP > Non-mus | Left Heschl's gyrus | Right PT | 357 | 0.02 | 69 | -24 | 21 |
| AP > Non-mus | Left Heschl's gyrus | Left PT | 140 | 0.01 | -66 | -15 | 9 |
| AP > Non-mus | Left Heschl's gyrus | Right preCG | 85 | 0.02 | 45 | 0 | 42 |
| AP > Non-mus | Left Heschl's gyrus | Right IFG,po | 28 | 0.04 | 48 | 15 | 21 |
| AP > Non-mus | Right Heschl's gyrus | Right aSMG | 50 | 0.04 | 69 | -15 | 27 |
| AP > Non-mus | Right Heschl's gyrus | Left pSTG | 19 | 0.04 | -63 | -18 | 3 |
| AP > Non-mus | Left PT | Right PT | 489 | 0.005 | 63 | -24 | 12 |
| AP > Non-mus | Left PT | Left PT | 322 | 0.002 | -63 | -21 | 6 |
| AP > Non-mus | Left PT | Right IFG,po | 103 | 0.02 | 48 | 15 | 21 |
| AP > Non-mus | Left PT | Right MTG | 71 | 0.02 | 45 | 0 | 42 |
| AP > Non-mus | Right PT | Left PT | 528 | 0.001 | -60 | -18 | 0 |
| AP > Non-mus | Right PT | Right PT | 355 | 0.005 | 60 | -12 | -3 |
| AP > Non-mus | Right PT | Right IFG,po | 264 | 0.008 | 45 | 15 | 24 |

Abbreviations: AP = absolute pitch musicians; aSMG = anterior supramarginal gyrus; IFG,po = inferior frontal gyrus, pars opercularis; MTG = middle temporal gyrus; Non-mus = non-musicians; $k$ = cluster size in voxels; $p_{FWE}$ = minimal family-wise error-corrected $p$ value in cluster; preCG = precentral gyrus; PT = planum temporale.

